# Single-cell transcriptome analysis reveals cell-cell communication and thyrocyte diversity in the zebrafish thyroid gland

**DOI:** 10.1101/2020.01.13.891630

**Authors:** Pierre Gillotay, Meghna Shankar, Benoit Haerlingen, Sema Elif Eski, Macarena Pozo-Morales, Inés Garteizgogeascoa Suñer, Susanne Reinhardt, Annekathrin Kränkel, Juliane Bläsche, Andreas Petzold, Nikolay Ninov, Gokul Kesavan, Christian Lange, Michael Brand, Vincent Detours, Sabine Costagliola, Sumeet Pal Singh

## Abstract

The thyroid gland regulates growth and metabolism via production of thyroid hormone in follicles composed of thyrocytes. So far, thyrocytes have been assumed to be a homogenous population. To uncover genetic heterogeneity in the thyrocyte population, and molecularly characterize the non-thyrocyte cells surrounding the follicle, we developed a single-cell transcriptome atlas of the zebrafish thyroid gland. The 6249-cell atlas includes profiles of thyrocytes, blood vessels, lymphatic vessels, immune cells and fibroblasts. Further, the thyrocytes could be split into two sub-populations with unique transcriptional signature, including differential expression of the transcription factor *pax2a*. To validate thyrocyte heterogeneity, we generated a CRISPR/Cas9-based *pax2a* knock-in line, which demonstrated specific *pax2a* expression in the thyrocytes. However, a population of *pax2a*-low mature thyrocytes interspersed within individual follicles could be distinguished, corroborating heterogeneity within the thyrocyte population. Our results identify and validate transcriptional differences within the nominally homogenous thyrocyte population.

**One-line summary:** Single-cell analysis uncovers latent heterogeneity in thyroid follicular cells.

**Graphical Abstract:** 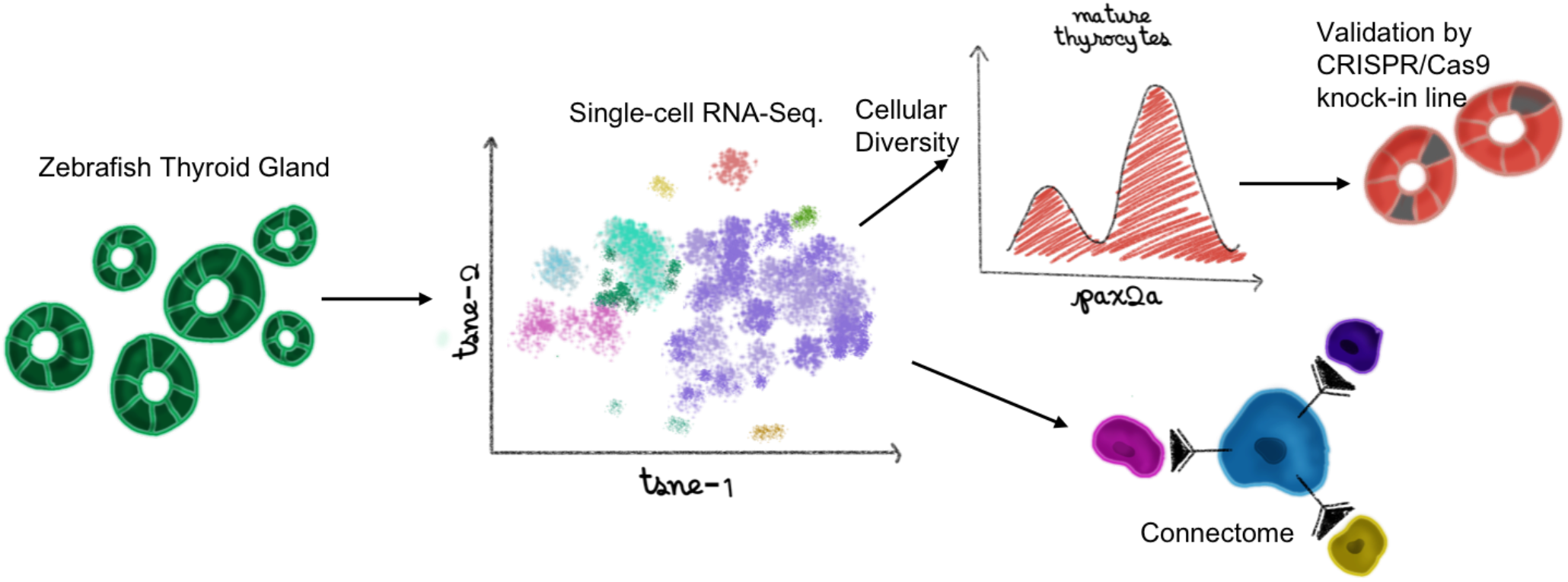

## Introduction

The thyroid gland produces hormones thyroxine (T4) and triiodothyronine (T3) that regulate body metabolism, growth, and development. Thyroid dysfunction, a disease afflicting almost 100 million people worldwide (1), is common and treatable by hormone replacement. If left untreated, however, it may result in profound adverse effects on the human body, including mental retardation, goiter or dwarfism.

The thyroid gland is an endocrine organ with an intricate structure enabling production, storage and release of the thyroid hormones. It contains numerous variable-sized spherical follicles composed of thyroid follicular epithelial cells, or thyrocytes. The thyrocytes generate the thyroid hormones in a multi-step process. They secrete and store thyroglobulin (TG) in the lumen of the follicles. Additionally, they intake iodide from the blood via sodium-iodide symporter (NIS / Slc5a5). At the interface between thyrocytes and the lumen, thyroid peroxidase (TPO) expressed by the cells catalyzes the coupling of iodide to tyrosyl residues of TG. Iodinated TG is absorbed back into the thyrocyte and cleaved by cysteine proteases in lysosomes to form T4 and T3 (2). Though the machinery responsible for the production of thyroid hormones by thyrocytes is well established, it remains unknown if all the thyrocytes resident in the thyroid gland are equally capable of generating thyroid hormones. In other words, the extent of molecular homogeneity between individual thyrocytes has not yet been investigated.

Additionally, the thyroid gland contains many cell-types with potential roles in modulating thyrocyte functionality. The gland contains an extensive distribution of blood vessels, which carry iodide to the thyrocytes and carry thyroid hormones away from them. The thyroid follicles are separated by a mesenchymal cell population, called connective tissue septa, which also divides the gland into lobules. The mammalian thyroid gland also contains parafollicular epithelial cells, or C-cells, that synthesize and secrete the hormone calcitonin. These parafollicular epithelial cells are, however, located outside the thyroid gland in fish and amphibians (3). Further, the presence of immune cells and innervation has been demonstrated within the thyroid gland (4, 5). Though we have a considerable understanding of these cell-types on a histological level, we still lack the molecular characterization of the thyroid gland cell ensemble. This extends to an incomplete appreciation of the impact of the diverse cell-populations on thyroid follicular cell physiology.

To uncover the diversity within the thyrocyte population, and further characterize the surrounding tissue at cellular resolution, we develop the first atlas of the thyroid gland at single-cell resolution. For this, we build on the progress in single-cell transcriptomics (6) to transcriptionally profile thousands of individual cells isolated from the thyroid gland of adolescent and adult zebrafish. We demonstrate that these profiles comprehensively represent the cells present in the zebrafish thyroid gland. Further, we demonstrate the segregation of thyrocytes into two transcriptionally distinct sub-populations. Utilizing the expression profiles of discrete cell populations, we build an intercellular signaling network to uncover communication between thyrocytes and the surrounding tissue. Finally, to enable easy access to the data, we have made the zebrafish thyroid gland atlas available for online browsing.

## Results

### Single-cell transcriptomics of the zebrafish thyroid gland

The zebrafish thyroid gland is composed of follicles scattered in the soft tissue surrounding the ventral aorta (Fig. 1 A, B). Ventral aorta extends from the outflow tract of the zebrafish heart and carries blood from the ventricle to the gills. Dissection of the ventral aorta associated region (detailed in Methods section) provided us with tissue that included the thyroid follicles and parts of zebrafish gills (Fig. 1C). Using *Tg(tg:nls-EGFP)* transgenic line, which labels thyrocytes with nuclear green fluorescence (Fig. 1D), we estimated presence of 5.9 ± 1.9 % thyrocytes within the dissociated region (Fig. 1E).

**Figure 1:**
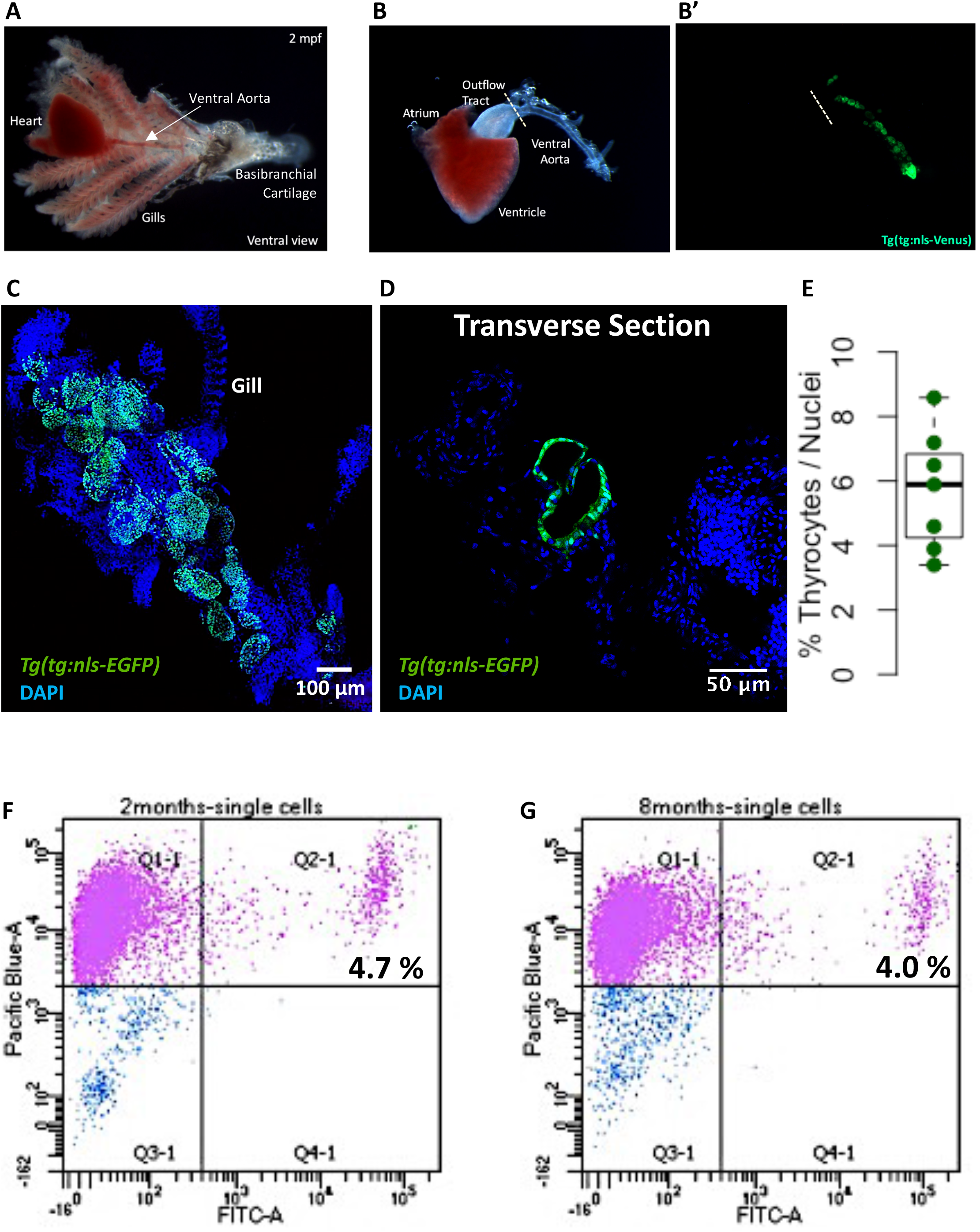
Isolation of zebrafish thyroid gland. **(A – B)** A brightfield image showing the zebrafish thyroid gland along with surrounding tissue. The thyroid follicles reside in the soft tissue surrounding the ventral aorta, which extends from the outflow tract of the heart into the gills towards the basibranchial cartilage in the lower jaw. The thyroid follicular cells, or thyrocytes, are labeled in green in the *Tg(tg:nls-mVenus-NTR)* transgenic line (B’). **(C)** Maximum intensity projection of 3D confocal stack obtained from the dissected thyroid gland labeled with DAPI. **(D)** Confocal scan of a transverse section across the dissected thyroid gland from *Tg(tg:nls-EGFP)* animal at 3 mpf. Sections were stained with DAPI to visualize cells surrounding thyroid follicles. **(E)** Boxplot depicting the proportion of thyrocytes present in transverse sections obtained from three *Tg(tg:nls-EGFP)* animals at 3 mpf. Each dot represents a transverse section. **(F – G)** Representative FACS plot of single cells from *Tg(tg:nls-mVenus-T2A-NTR)* animals at 2 mpf (F) and 8 mpf (G). Calcein (Pacific Blue) labels live cells, while green fluorescence (FITC) labels thyrocytes. Percentage values represent proportion of calcein+ thyrocytes within total calcein+ cells.

To generate the molecular catalogue of the thyroid gland at cellular resolution, we sampled the organ from two ages of zebrafish: 2 month post-fertilization (mpf) and 8 mpf (Supp. Figure 1). The time points span adolescent to adult transition in zebrafish, with animals containing fully differentiated functional organs at both stages. By 2 mpf, the adolescent animals have completed morphogenesis, but are yet to reach sexual maturity. The animals sampled at 2 mpf were on average 2.6 cm in length and 123.8 mg in weight. In contrast, fish at 8 mpf are sexually mature adults, with an average length of 3.5 cm and an average weight of 294.4 mg (Supp. Figure 1). To characterize the organ cell-types in an unbiased manner, we dissected out the entire thyroid gland (Fig. 1B, C) from six animals at each stage, and prepared the single-cell suspension for cDNA library preparation. To guide thyroid gland dissection, we utilized the *Tg(tg:nls-mVenus-T2A-NTR)* zebrafish reporter line (7) that labels thyrocytes with bright yellow fluorescent protein (Fig. 1B’). The micro-dissected tissue was dissociated using enzymatic digestion. The single-cell suspension was stained with calcein, which specifically labels live cells with blue fluorescence. The live cells were then enriched using FACS (Fig. 1F - G) to limit false positive signals from dead and/or ruptured cells (8). Thyrocytes consisted of around 4 % of the alive cells at both stages, comparable to the percentage obtained by immunofluorescence analysis (Fig. 1E). Twelve thousand live-cells, pooled from six animals, were collected in separate tubes according to age and profiled using droplet-based high-throughput single-cell RNA-sequencing provided by 10X Genomics (9, 10). Droplet-based methods encapsulate cells with single-Poisson distribution (10). This leads to approximately 50% cell capture rate, which is the ratio of the number of cells detected by sequencing and the number of cells loaded. The 10X Genomics pipeline uses molecule and cell-specific barcoding allowing transcript quantification without amplification bias (11, 12). Using the Cell Ranger bioinformatics pipelines, the resulting Next-Generation Sequencing libraries were mapped to the zebrafish genome, de-multiplexed according to their cellular barcodes and quantified to generate gene/cell UMI (unique molecular identifier) count tables. The Cell Ranger pipeline provided us with 13,106 sequenced cells from 24,000 input cells (54.6 % cell capture rate). Quality-based exclusion of single-cell transcriptomes was implemented based on mean library size, percentage of mitochondrial reads and number of genes detected per cell. On average, we detected 6,012 UMIs and 1,303 genes per cell (Supp. Figure 2). The process recovered in total 6,249 cells out of 13,106 sequenced cells (47.7 % retention rate), providing single-cell transcriptomic profiles for 2986 and 3263 individual cells for 2 mpf and 8 mpf, respectively.

### Identification of cell-types present in the zebrafish thyroid gland

To aid with visualization of the zebrafish thyroid gland single-cell RNA-Seq (scRNA-Seq) data, we projected the cellular profiles onto t-distributed stochastic neighbor embedding (t-SNE) plots, a non-linear dimensionality reduction technique (13) (Fig. 2A). Using unsupervised graph-based clustering, we identified seven clusters for the thyroid gland. Using the expression of genes involved in thyroid hormone production, we could identify one of the clusters as thyroid follicular cells (Fig. 2B – D). Specifically, the cluster displayed high relative expression of *tg* gene, which was further enriched by background correction (Supp. Fig. 3); thereby demonstrating that the cells represented differentiated thyroid follicular cells. The cluster, labeled as thyrocytes, contains 267 cells. This represents 4.2 % of the total cells recovered after quality control, similar to the proportion of thyrocytes quantified in the dissociated tissue by imaging and FACS (Fig. 1E – G), suggesting lack of thyrocyte loss during the sequencing procedure.

**Figure 2:**
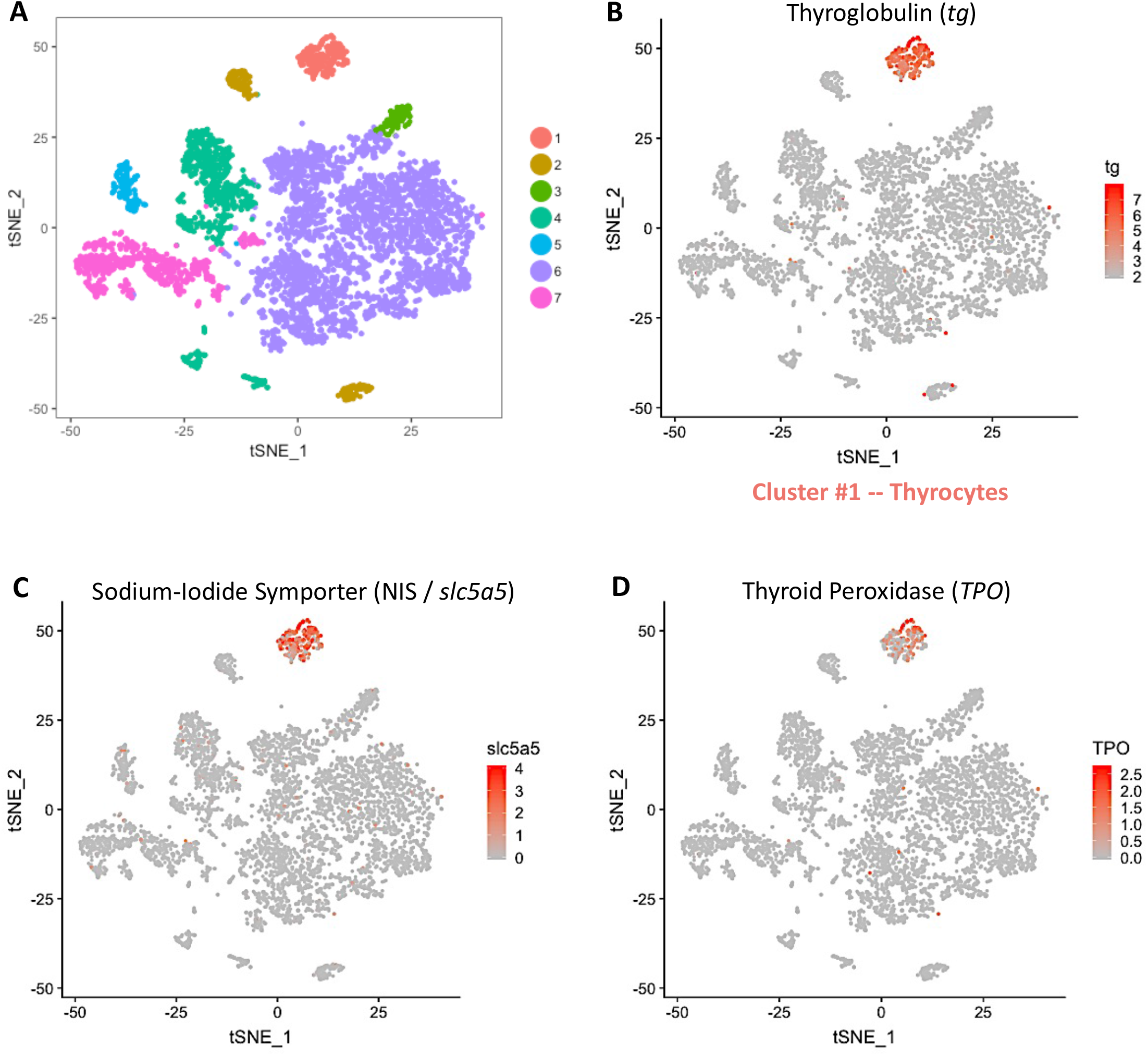
Single-cell RNA-Seq. of the zebrafish thyroid gland. **(A)** A t-SNE plot displaying the 6249 single-cells profiled in the zebrafish thyroid gland atlas. The colors represent cell clusters denoting a specific cell-type. **(D – F)** Cluster #1 represents the thyrocytes that express *tg*, *slc5a5* (NIS) and *tpo*.

To define the identity of the remaining cell clusters, we generated cluster-specific marker genes by performing differential gene expression analysis (Fig. 3A) (Supp. Table 1). For four clusters, the marker genes included one or more known cell type–specific identifiers. This included *gpr182* for endothelial cells; *acta2* for musculature; *fcer1gl* for immune cells; and *ponzr3* for cells from zebrafish gills (Fig. 3B – E). Based on these cell identifiers, the atlas includes 233 endothelial cells, 135 muscle lineage cells, 914 immune cells and 199 cells from zebrafish gills. Notably, the endothelial cell cluster includes blood vessels (*flt1* and *kdrl*) and lymphatic vessels (*mrc1a*, *prox1a*, *flt4* and *lyve1b*) (Supp. Fig. 4); while the immune cell cluster includes macrophages (*mpeg1.1* and *mfap4*), neutrophils (*lyz*) and lymphocytes (*il4*, *il13* and *il11b*) (Supp. Fig. 5).

**Figure 3:**
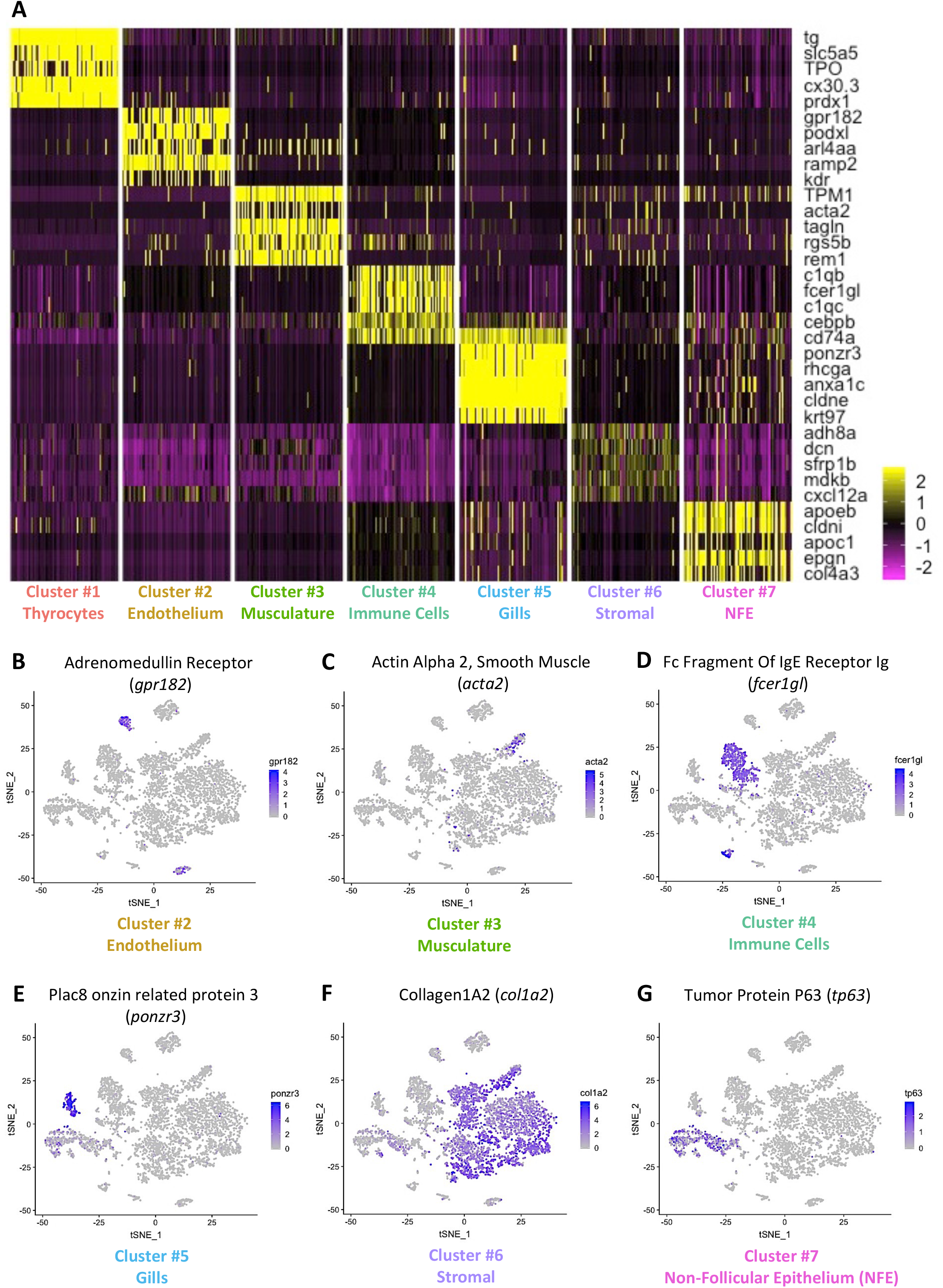
Gene expression signature of the different cell-types in the zebrafish thyroid gland. **(A)** Heatmap depicting five genes specifically expressed in each one of the seven clusters of the zebrafish thyroid gland atlas. **(B – G)** t-SNE plots overlaid with the expression of a gene specific to each of the cluster. The endothelium cluster (cluster #2) is a mix of blood vessels and lymphatic vessels (see Supp. Fig. 4), while the immune cell cluster (cluster #4) is a mix of macrophages, neutrophils and lymphocytes (see Supp. Fig. 5).

For the remaining two clusters (number six and seven), we identified marker genes that hinted towards identity of the cell-type. Specifically, *col1a2* and *tp63* enriched in cluster number six and seven respectively (Fig. 3 F – G), are known markers of fibroblasts (14, 15) and epithelial tissue (16–18). We performed gene-ontology (GO) enrichment analysis of the marker genes to aid with classification (Supp. Fig. 6). Cluster six demonstrated an enrichment of ‘extracellular matrix structural constituent’, ‘connective tissue development’ and ‘extracellular space’, confirming the presence of tissue fibroblasts in this cluster. Thus, we labelled cluster six as ‘Stromal’ cells. Cluster seven displayed an enrichment of ‘cell motility’, ‘cell migration’ and ‘epithelium development’, suggestive of epithelial cells. Hence, we labelled cluster seven as ‘Non-Follicular Epithelium (NFE)’, to distinguish them from the thyroid follicular epithelial cells. Our data contains 3670 stromal cells and 831 non-follicular epithelial cells.

We validated the presence of blood vessels, macrophages and stromal cells in the thyroid gland using tissue specific transgenic lines (Fig. 4A – C).

**Figure 4:**
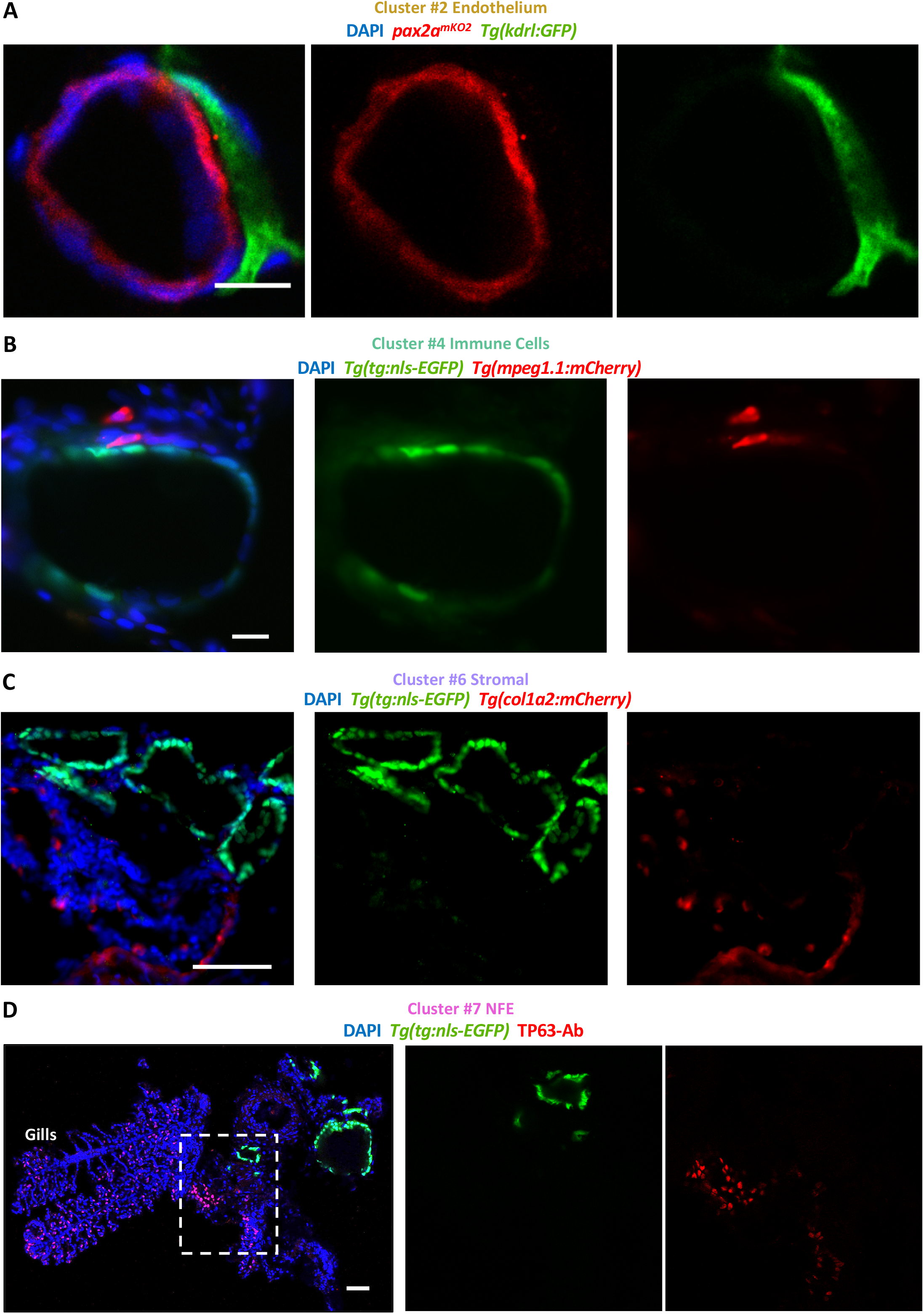
Immunofluorescence-based visualization of cell-types surrounding zebrafish thyroid follicles. Images show immunofluorescence labeling of thyroid gland from adult zebrafish. Transverse sections were utilized for imaging. The organ was isolated from tissue-specific transgenic lines to allow marking of a particular cell-type adjacent to the thyroid follicle. Blood vessels were marked using *Tg(kdrl:EGFP)* **(A)**, macrophages using *Tg(mpeg1.1:mCherry)* **(B)** and stroma using *Tg(col1a2:mCherry)* **(C)**. Thyrocytes were labeled with *pax2a*^*mKO2*^ expression in (A) (described in Fig. 7) and *Tg(tg:nls-EGFP)* expression in (B – C). **(D)** NFE was labeled using antibody against TP63 in sections of the thyroid gland isolated from *Tg(tg:nls-EGFP)* animals. Gills are marked based on their morphological appearance. DAPI labels nuclei. Scale bars: 10 μm (A - B), 50 μm (C – D).

Immunofluorescence (IF) analysis demonstrated physical proximity between thyrocytes and blood vessels (Fig. 4A). Notably, we observed a subset of macrophages in direct contact with thyroid follicles (Fig. 4B). In addition, we visualized NFE by immunostaining against TP63 antibody (Fig. 4D), which revealed NFE scattered throughout the gills and in the region adjacent to the follicles. Thus, the IF analysis successfully confirmed the presence of different cell types identified in the single-cell atlas.

Our marker gene identification further established additional genes enriched in a single cell-type in the thyroid gland (Fig. 3A) (Supp. Table 1). For instance, we identified *cx30.3*, a connexin gene and *prdx1*, a gene involved in the antioxidant response, to be specifically expressed in the thyrocytes. To enable further investigation of the clusters and gene expression profiles, we have developed an interactive webtool for online browsing (https://sumeet.shinyapps.io/zfthyroid/).

### Development of autocrine and paracrine signaling networks in the thyroid gland using known ligand-receptor interactions

Having defined the cell types of the thyroid gland, we quantified potential cell-cell interactions between thyrocytes and all cell types present in the organ (Fig. 5A) based on a reference list of approximately 3,100 literature-supported interactions containing receptors and ligands from receptor tyrosine kinase (RTK), extracellular matrix (ECM)-integrin, chemokine and cytokine families (19). Although anatomical barriers between cell types are not modeled in this analysis, we restricted the analysis to secreted ligands for NFE, stroma and gills -- cell types that are physically separated from thyrocytes (Fig. 4C - D). For the remaining cell types, secreted and cell-membrane tethered ligands were considered. The expression patterns of ligand-receptor pairs revealed a dense intercellular communication network (Fig. 5B). The network consisted of 272 ligands expressed on different cell-types with a corresponding receptor expressed on the thyrocytes (Supp. Table 2). For instance, the stromal cells express the ligand *lpl* (Lipoprotein Lipase) that signals through the *lrp2a* (zebrafish homologue of Megalin) receptor (Fig. 5C). Stromal and smooth muscle cells express *dcn* (Decorin) whose receptor *met* is expressed by thyrocytes. Further, the ligand *cyr61* is broadly expressed in the thyroid gland, with one of its receptors, *itgb5*, an integrin isoform, expressed specifically by the thyrocytes. The identified interactions also include autocrine signaling. For example, the ligand *sema3b* and its receptor *nrp2a* are both present on thyrocytes. GO-analysis for identified ligand-receptor pairs revealed genes involved in ‘PI3K-Akt signaling pathway’, ‘MET signaling’ and ‘integrin binding’ (Supp. Fig. 7).

**Figure 5:**
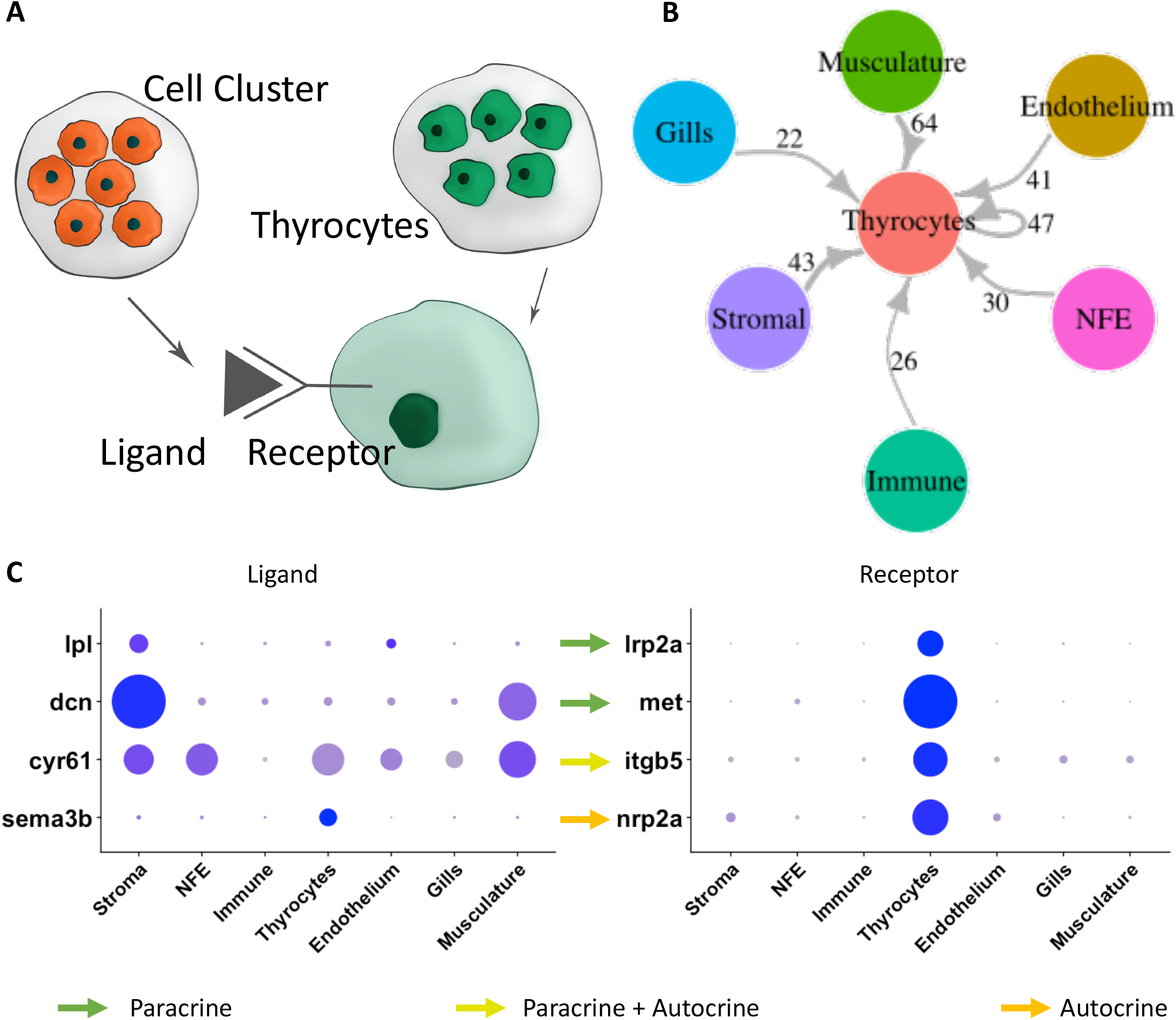
Connectome of the zebrafish thyroid gland identifies a dense intercellular signaling network. **(A)** To build a connectome for the atlas, the ligands expressed specifically in each cell-type were matched with their corresponding receptors in the thyrocytes. **(B)** A highly connected intercellular interaction network is identified by the connectome. The number of ligand-receptor pairs identified between two cell-types is denoted alongside the arrows. For NFE, Gills and Stromal cells, the connectome was restricted to secreted ligands. **(C)** A dotplot depicting examples of paracrine and autocrine signaling in the thyroid gland. The dots represent expression level in the different cell-types of the atlas.

### Thyrocytes are composed of transcriptionally distinct sub-populations

Next, we characterized the transcriptional differences within the thyrocyte population. For this, we bioinformatically isolated the thyrocytes, and re-performed the clustering pipeline on the isolated cell population. With this, we could segregate the thyrocytes into two smaller clusters (Fig. 6A), labeled as ‘Cluster_Blue’ and ‘Cluster_Red’. The two clusters displayed differences in the expression levels of 265 genes (Fig. 6B) (Supp. Table 3). Notably, Cathepsin B (*ctsba*) is significantly downregulated in the blue cluster (Fold change = 1.6, p-value = 1.47×10^−9^) (Fig. 6B – C). Cathepsin B is a cysteine protease that is involved in the processing of iodinated thyroglobulin to T4 and T3 in the thyrocyte lysosomes (2, 20). Moreover, fusion of Cathepsin B and EGFP has been previously used to track thyroid hormone processing lysosomes in rat thyroid epithelial cell lines (21).

**Figure 6:**
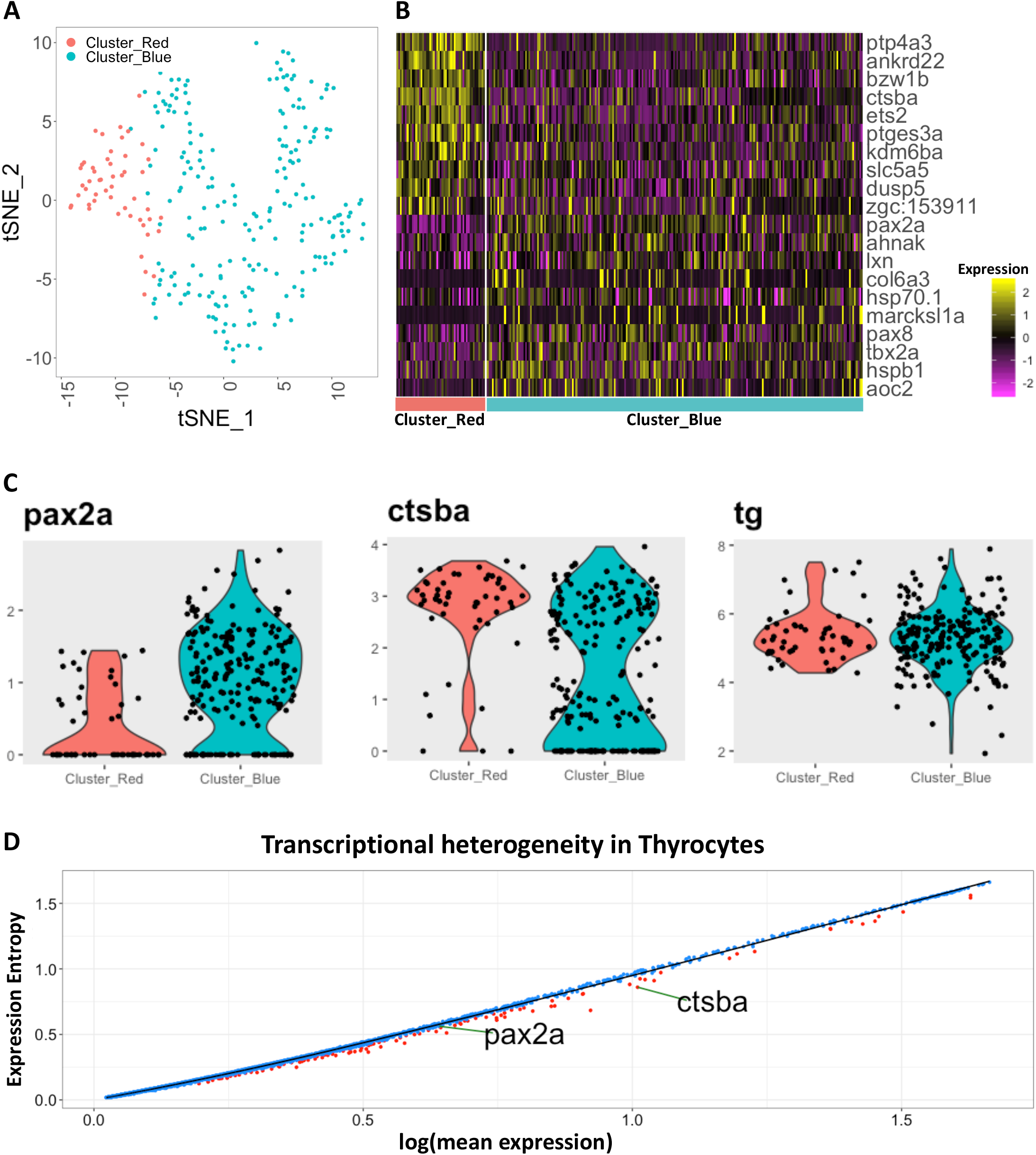
Thyrocytes can be subdivided into two transcriptionally distinct sub-populations. **(A)** Unsupervised clustering of the thyrocyte population identifies two sub-populations. **(B)** Heatmap depicting the top ten most differentially expressed genes between the two sub-populations. **(C)** Violin plots depicting the expression levels of *pax2a*, *ctsba* and *tg* in each sub-population. Y-axis represents scaled data. **(D)** Dot plot depicting expression entropy on Y-axis against average gene expression on X-axis for the thyrocyte population. Each dot depicts a gene, with red dots depicting genes that show statistically significant (p-value < 0.05) difference in entropy from expected value. Expected value is represented by black regression line. *pax2a* and *ctsba* are marked on the graph.

Along with higher expression of Cathepsin B, the red cluster displayed significant downregulation of *pax2a* expression (Fold change = 1.7, p-value = 8.24×10^−9^) (Fig. 6B-C). *pax2a* belongs to the PAX (paired box DNA-binding) domain containing family of transcription factors. The loss of *pax2a* expression in the red cluster is notable, as *pax2a* is an important regulator of thyrocyte development (22). Zebrafish thyroid primordium expresses *pax2a* at 24 hpf (22), which is required for specification of the thyroid follicles (23, 24). Consequently, zebrafish lacking *pax2a* fail to develop thyroid follicles (22), which is similar to the Pax8 knock-out phenotype in mouse (25). The low expression of *pax2a* in the red cluster, without a difference in *tg* expression (Supp. Fig. 8), suggests the presence of a thyrocyte sub-population with a distinct gene expression signature.

Independent analysis of genetic entropy, a measure of the degree of uncertainty, revealed transcriptional heterogeneity in 231 genes in the thyrocyte population (Fig. 6D) (Supp. Table 4). Genes displaying statistically significant entropy (p-value < 0.05) included *pax2a* and *ctsba*, corroborating their expression heterogeneity within thyrocytes.

### Generation of *pax2a* knock-in reporter line

To validate the heterogeneity among the zebrafish thyrocytes, we focused on the expression of *pax2a* transcription factor. We generated a knock-in line by inserting monomeric Kusabira Orange 2 (mKO2) fluorescent protein to the 3’ end of the endogenous *pax2a* genomic location (Fig. 7A). The *pax2a*^*pax2a-T2A-mKO2*^ (abbreviated as *pax2a*^*mKO2*^) reporter expression overlapped with PAX2A antibody staining in a majority of regions at 9.5 hours post-fertilization (Fig. 7B). Moreover, the knock-in line displayed mKO2 fluorescence in the otic vesicle, mid-hindbrain boundary, optic stalk, pronephros and the thyroid gland (Fig. 7C – F, Supp. Movie 1), mimicking known expression of *pax2a* during zebrafish development (26). Additionally, in order to assess whether the dynamics of mKO2 expression would follow modifications in the expression of endogenous *pax2a,* we used CRISPR/Cas9 technology to generate F0 knock-outs (also known as Crispant (27)) of *pax2a* gene in our *pax2a*^mKO2^ line. The crispants displayed defects in thyroid morphogenesis (Fig. 7G – H), mimicking the phenotype of *pax2a* loss-of-function mutation (22). Live imaging of crispants at 55 hpf revealed strong decrease of mKO2 expression (Fig. 7G – H), thereby corroborating the faithful recapitulation of *pax2a* expression by the newly generated reporter line.

**Figure 7:**
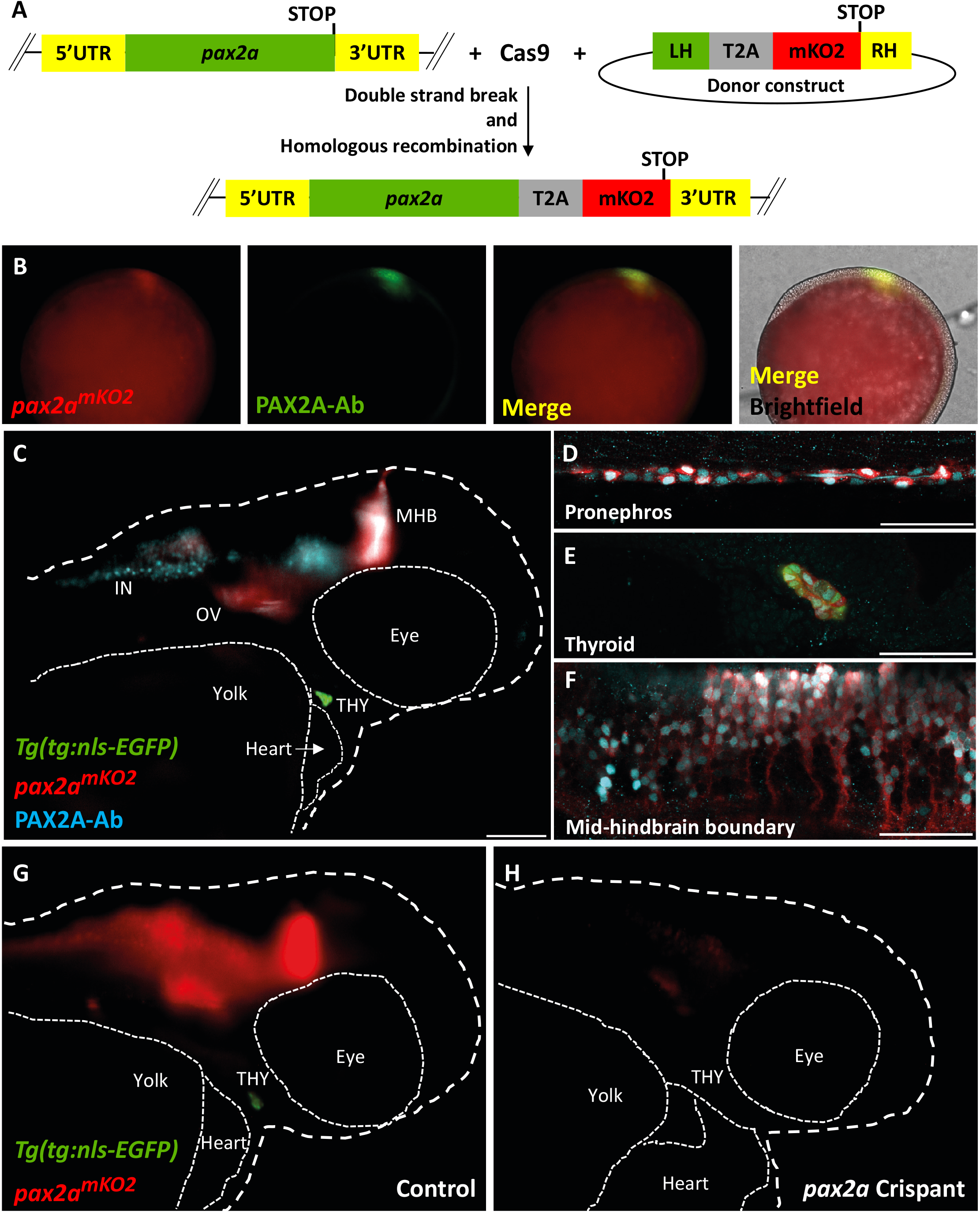
*pax2a*^*mKO2*^ knock-in line faithfully reports *pax2a* expression and knock-down. **(A)** Schematic of the knock-in strategy used to generate the *pax2a*^*mKO2*^ line. Double strand break was induced between the penultimate codon and the STOP codon of *pax2a* gene using CRISPR/Cas9. DNA repair integrates the donor construct at the site of double strand break, resulting in a *pax2a* reporter line. The donor construct contains T2A-mKO2 reporter cassette flanked by left homology (LH) and right homology (RH) arms. **(B)** Whole mount immunofluorescence of 9.5hpf *pax2a*^mKO2^ embryos stained with anti-mKO2 antibody (red) and anti-PAX2A antibody (green). Anterior is to the left, and dorsal side is to the top. **(C)** Whole mount immunofluorescence of 55 hpf *pax2a*^*mKO2*^; *Tg(tg:nls-EGFP)* stained with PAX2A antibody (PAX2A-Ab) displays an overlap of mKO2 and PAX2A-Ab signal. The otic vesicle (OV), mid-hindbrain barrier (MHB), interneurons (IN) and thyroid gland (THY) is labelled. **(D – F)** Confocal microscopy imaging of a sagittal section of a 55 hpf *pax2a*^*mKO2*^; *Tg(tg:nls-EGFP)* embryos showing co-localization of mKO2 and *pax2a* in the pronephros (D), thyroid gland (E) and mid-hindbrain barrier (F). In the thyroid gland, mKO2, PAX2A-Ab and thyrocyte-specific GFP (green) show co-localization. Scale bars: 100μm (C) and 50μm (D – F). Anterior to the right, white dashed line represents the outline of the embryo. **(G – H)** Snapshots from live imaging of 55 hpf *pax2a*^mKO2^; *Tg(tg:nls-EGFP)* embryos injected with sgRNA targeting *pax2a* coding sequence. The anterior part of a representative control embryo (G) is shown alongside a representative crispant (H). Crispants display a strong reduction of mKO2 fluorescence, as well as an absence of GFP signal suggesting absence of thyroid (THY) tissue.

### Segregation of thyrocyte sub-populations based on *pax2a* reporter expression

Upon investigating the fluorescence expression of the *pax2a* reporter in the thyroid gland of adult zebrafish, we found strong and specific expression of *pax2a* reporter in the thyrocytes lining the thyroid follicles (Fig. 8A – D). Although a majority of thyrocytes displayed uniform expression of *pax2a* reporter, we could identify a small population of *pax2a*^*mKO2*^-Low thyrocytes (Fig. 8B – D). The *pax2a*^*mKO2*^-Low thyrocytes were not segregated, but scattered throughout the gland, thereby suggesting a mixing of the two thyrocyte sub-populations.

**Figure 8:**
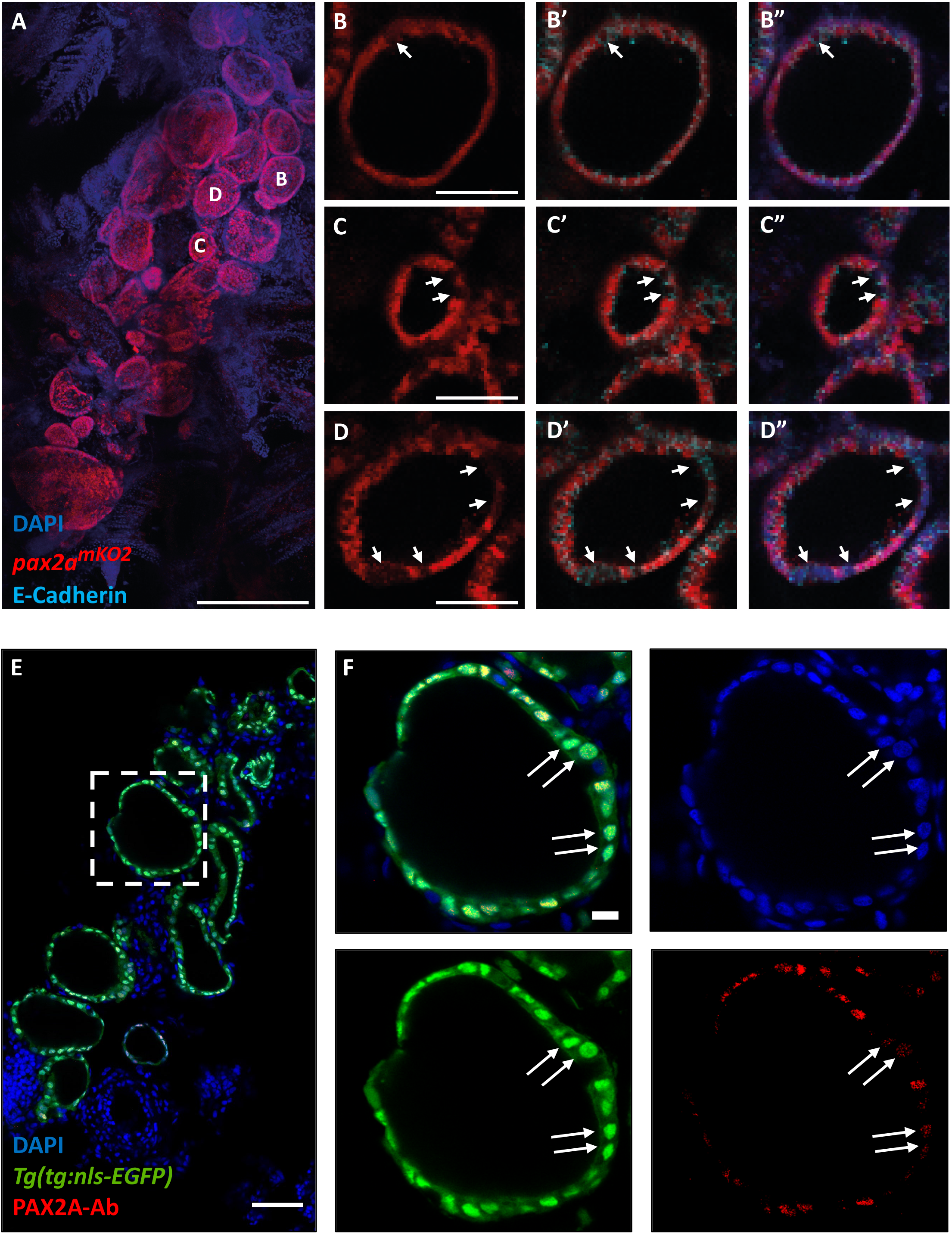
Immunofluorescence-based validation of thyrocyte heterogeneity. **(A-D)** Analysis of 3 mpf thyroid gland from *pax2a*^mKO2^ zebrafish shows heterogeneity in *pax2a* reporter expression. **(A)** Whole mount confocal imaging of a 3 mpf *pax2a*^mKO2^ thyroid labelled with mKO2 (red), E-cadherin (cyan, not shown in ‘A’ for clarity reasons) and DAPI (dark blue) for nuclear localisation. **(B – D)** Optical sections of three follicles, with mKO2-Low cells labelled with arrows. E-cadherin (B’ – D’) and DAPI (B’’ – D’’) staining shows that absence of mKO2 signal does not correspond to an absence of cells. Anterior to the bottom of the pictures. **(E)** Confocal image of thyroid gland section from *Tg(tg:nls-EGFP)* at 4 mpf stained with PAX2A antibody and DAPI. The dotted region is displayed at high magnification in **(F)**. Arrows marks thyrocytes displaying low PAX2A staining. Notably, PAX2A-Low thyrocytes display *tg*-driven EGFP expression, demonstrating their differentiated status. Scale bars: 250 μm (A), 50 μm (B – E), 10 μm (F).

To validate pax2a expression heterogeneity at a protein level, we performed immunostaining against PAX2A in thyroid gland obtained from *Tg(tg:nls-EGFP)* animals (Fig. 8E). For antibody staining, we utilized 8 μm thin sections of the thyroid gland to ensure uniform antibody penetration to all cells. Confocal imaging of the stained sections demonstrated the presence of PAX2A-Low and PAX2A-High thyrocytes (Fig. 8F). Notably, both PAX2A-Low and -High cells display *tg* promoter-driven EGFP expression, thereby confirming their differentiated status.

Further, to quantify the proportions of *pax2a*^*mKO2*^-Low and -High thyrocytes, we performed FACS analysis on *pax2a*^*mKO2*^; *Tg(tg:nls-EGFP)* double transgenic line (Fig. 9A – C). The *Tg(tg:nls-EGFP)* zebrafish line labels the thyrocyte population in green fluorescence (27). We restricted our analysis to the thyrocyte population by gating for GFP+ cells in the thyroid gland (Fig. 9A). Within the thyrocyte population, the cells displayed a normal distribution of GFP fluorescence; however, thyrocytes could be split into two sub-populations based on the levels of *pax2a* reporter expression (Fig. 9B – C). Specifically, 75% of thyrocytes (202 out of 268 cells) displayed *pax2a*^*mKO2*^-High fluorescence, while 25% of thyrocytes (66 out of 268 cells) displayed *pax2a*^*mKO2*^-Low fluorescence levels.

**Figure 9:**
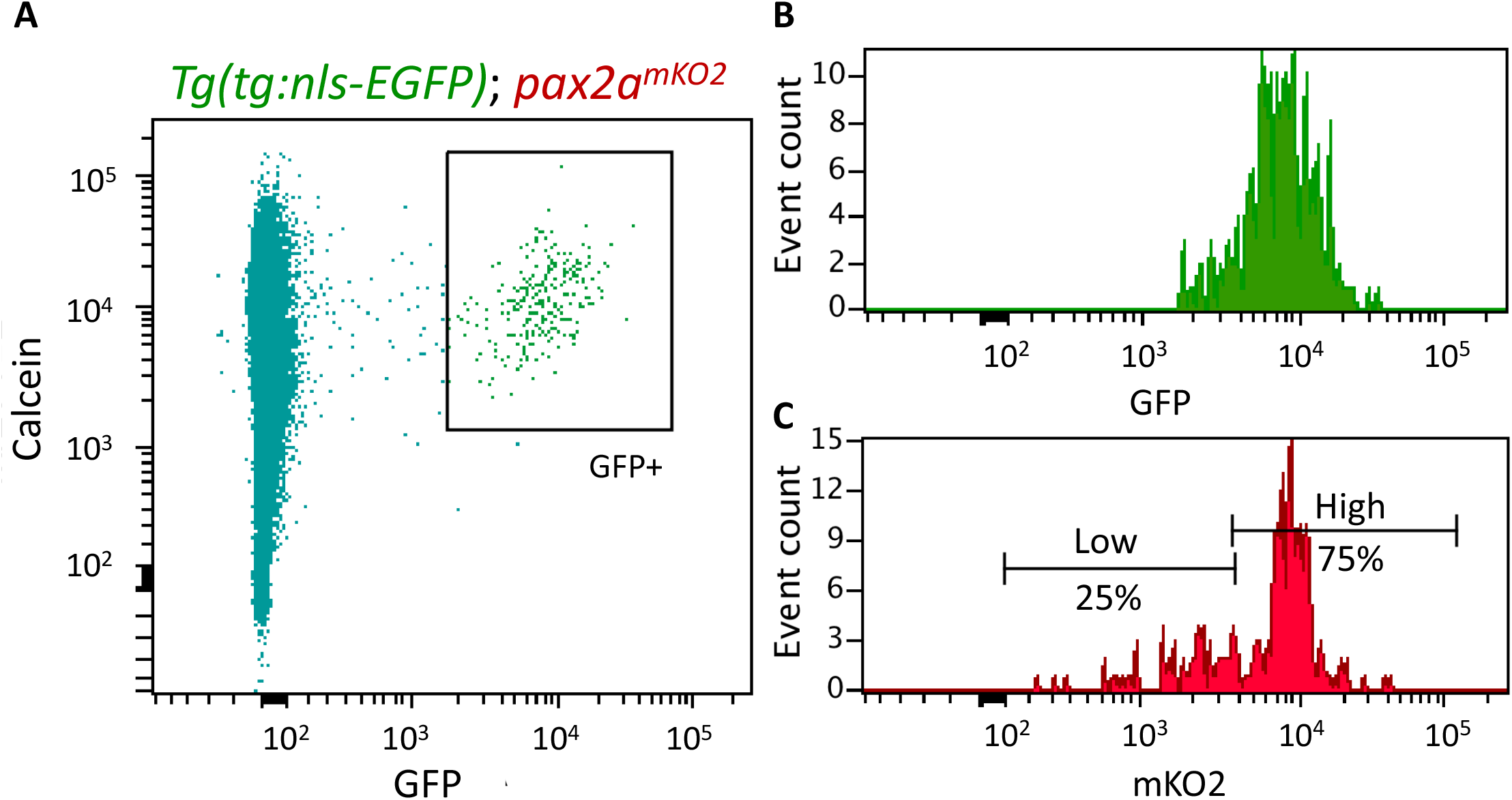
Flow cytometry-based validation of thyrocyte heterogeneity. Cells from the thyroid gland of 5 mpf *Tg(tg:nls-GFP)*; *pax2a*^*mKO2*^ animals were labelled with calcein (live cell marker) and analysed using FACS. **(A)** A FACS plot showing calcein on X-axis and GFP on Y-axis. The box encompassing the GFP+ cells represents the thyrocyte population, which was gated for further analysis. **(B)** Histogram showing the distribution of GFP intensity in thyrocytes. **(C)** Histogram showing the distribution of mKO2 intensity in thyrocytes. Thyrocytes were selected by gating for GFP+ population. Horizontal lines indicate the mKO2-Low and mKO2-High expression level, with percentage values representing proportion of thyrocytes with mKO2-Low and mKO2-High expression.

In summary, the analysis of *pax2a* knock-in line, along with PAX2A immunofluorescence imaging, validates the identification of thyrocyte sub-populations within our single-cell RNA-Seq. data, and clearly demonstrates, for the first time, the presence of transcriptionally diverse sub-populations of thyrocytes present in the thyroid gland.

## Discussion

We have applied for the first time unbiased single-cell gene expression analysis to the thyroid gland. In contrast with the mainstream view that thyrocytes constitute a molecularly uniform population, we identify two transcriptionally distinct sub-populations of thyrocytes. The two sub-populations differed, among other genes (Supp. Table 3), in the expression levels of a transcription factor *pax2a* and a cysteine protease Cathepsin B (*ctsba*) (Fig. 6B – C). Cathepsin B is particularly notable as it enables the liberation of thyroid hormone from thyrocytes by proteolytic processing of thyroglobulin (2, 20).

We validate the heterogeneity among the thyrocytes using a newly generated knock-in reporter line for *pax2a* gene (Fig. 7). The knock-in reporter line was generated using CRISPR/Cas9-based insertion of mKO2 fluorescent protein in the endogenous *pax2a* genomic location. The *pax2a* knock-in line faithfully recapitulates the embryonic expression of *pax2a* gene (Supp. Movie 1, Fig. 7B – F). Using the *pax2a* reporter line to characterize the adult thyroid gland, we demonstrate the presence of *pax2a*^*mKO2*^-Low thyrocytes in the follicles (Fig. 8). Notably, *pax2a*^*mKO2*^-Low and *pax2a*^*mKO2*^-High thyrocytes are present in the same follicle (Fig. 8C – D, F), raising the possibility of contact-mediated interactions between the two sub-populations. It would be of interest to build on this study and investigate the functional and replicative differences among the two sub-populations of thyrocytes.

Our single-cell transcriptomics atlas provides a comprehensive genomics resource to study the zebrafish thyroid gland in unprecedented detail. We performed unbiased profiling of the thyroid gland, without enrichment for a specific cell-type. This allowed us to capture yet poorly characterized cell-populations within the thyroid gland. Specifically, we provide the molecular characteristics of the stromal tissue present in the zebrafish thyroid gland. The stromal cells (Fig. 4C) display enrichment of extra-cellular matrix (ECM) related genes (Supp. Table 1) and are possibly homologous to the mesenchymal connective septa found in the mammalian thyroid gland. The connective septa helps cluster the thyroid follicles into lobules. Notably, the expression of fgf ligands from the mesenchymal septa cells has been implicated in lobe formation during mouse thyroid gland development (28). It would be of interest to test if similar morphological clustering of the thyroid follicles exists in zebrafish, and the role the stromal cells play during development and growth of the thyrocytes.

Our atlas further identifies a non-follicular epithelial (NFE) cell-population present near the zebrafish thyroid follicles. A subset of NFE are present in the gills (Fig. 4D) and may potentially represent a progenitor population for the gills, similar to the TP63+ basal layer in the zebrafish (29) and mammalian (30) epithelium. We also observe NFE outside the gills (Fig. 4D), which may play a different role. It is interesting to note that epithelial cells apart from follicular and parafollicular cells have been observed in the mammalian thyroid gland. In a report from Dr. E. Baber published in 1876 (31), histological examination of the dog thyroid gland displayed the presence of cells “beside the stroma, lymphatics, blood vessels, & cells between the vesicles”. Dr. Baber labeled the cells as ‘parenchyma’, and noted the existence of “numerous cells differing markedly in size and shape from the epithelial cells amongst which they lie” (31). In 1907, Dr. Sophia Getzowa described an epithelial cells containing structure called the Solid Cell Nests (SCN) of the thyroid (32). SCN are lumen containing irregular structures located within the thyroid in mammals (33). SCN contain two types of epithelial cells: main cells and C-cells, expressing TP63 and calcitonin respectively (34). Notably, the NFE cells we identified in the zebrafish thyroid gland are marked with TP63 expression (Fig. 3G, 4D), raising the possibility of their homology with the main cells of the SCN. C-cells, however, exist in the ultimobranchial bodies, which lies outside the thyroid gland in zebrafish. The ultimobranchial bodies are the zebrafish homologues of parafollicular cells and are located as a pair of follicles on top the sinus venous, adjacent to the atrium and oesophagus (3). Cells adjacent to the atrium were removed during our dissections (Fig. 1B). Additionally, NFE cells identified in our atlas do not express the zebrafish homologue of calcitonin (*calca*) (Supp. Table 1), and thus it is unlikely that the NFE cells we have identified would be related to cells of parafollicular origin. Currently, the developmental origin of NFE cells and their role in thyroid gland is unclear. To study the cell-population, transgenic zebrafish reagents driving expression using the *tp63* regulatory region (35) could be utilized in future.

To survey the communication between thyrocytes, the functional unit of the thyroid gland, and the other cell-types present in the thyroid gland, we constructed a cellular interaction network (Fig. 5B). The network was built by matching the expression of ligands in the diverse cell-types with the expression of receptor in the thyrocytes (Supp. Table 2) (19). Based on literature survey, we manually identified multiple interacting genes that have been implicated in thyroid diseases. For instance, the ligand Decorin (*dcn*) is expressed by the stromal cells and musculature, with its receptor, MET, present on thyrocytes (Fig. 5C). Decorin, a secreted proteoglycan, is considered a “guardian from the matrix” (36), as it is an antagonist of growth factor signaling. Importantly, Decorin expression has been reported to be downregulated in thyroid cancer samples (37). Thus, stromal cells could modulate Decorin to control thyrocytes growth. Further, interactions for CYR61 (associated with Graves’ Disease (38)), LRP2 / Megalin (involved in thyroglobulin processing (39)) and NRP2 (associated with thyroid cancer metastasis (40)) were identified (Fig. 3C). The hypothesis generated by the theoretical ligand-receptor interaction network can be tested *in vivo* in zebrafish or *in vitro* by manipulation of thyrocytes in thyroid organoid models (41) to gain valuable insight into thyroid gland homeostasis.

The current atlas is restricted to healthy adolescent and adult thyroid gland. The two stages represent a period of slow growth in zebrafish. Thus, genes driving cellular proliferation might be repressed at these stages. Additionally, the low number of cells per cluster obtained at each stage restricts an in-depth analysis of the transcriptional difference with age. In future, it would be of interest to extend the atlas by increasing cell numbers and by including single-cell transcriptomics from embryonic and old fish, providing a comprehensive resource for development, homeostasis and aging of the thyroid gland. It would be of further interest to profile zebrafish models of thyroid disorder (42, 43) to understand the cellular and molecular changes underlying organ dysfunction. Combined with the power of CRISPR/Cas9 based screen that we have established for the thyroid gland (27), this resource will provide a roadmap for the functional elucidation of cell type specific programs during thyroid gland growth and homeostasis.

In summary, our work provides the first molecular map of the zebrafish thyroid gland at cellular resolution. The atlas contains the molecular characterization of the thyroid gland stromal population, identification of non-follicular epithelial cells, and demonstrate the transcriptional heterogeneity among zebrafish thyrocytes. Further, by constructing cell-cell communication network, the atlas provides clues into tissue dynamics present within the organ. Finally, the dataset has been made available for browsing via an interactive webtool (https://sumeet.shinyapps.io/zfthyroid/). We hope that our efforts will expand the understanding of thyrocytes beyond a nominally homogenous endocrine cell population; providing a complex picture of the diversity in thyrocyte identity and function.

## Methods

### Zebrafish strains and husbandry

Wild-type or transgenic zebrafish of the outbred AB, WIK, or a hybrid WIK/AB strain were used in all experiments. Zebrafish were raised under standard conditions at 28 °C. Animals were chosen at random for all experiments. Published transgenic strains used in this study were *Tg(tg:nls-mVenus-T2A-NTR)* (7), *Tg(tg:nls-EGFP)* (27), *Tg(kdrl:GFP)*^la116^ (44), *Tg(mpeg1.1:mCherry)*^gl23^ (45) and *Tg(col1a2:LOXP-mCherry-NTR)*^*cn11*^ (referred as *Tg(Col1a2:mCherry)*) (14). Experiments with *Tg(tg:nls-mVenus-T2A-NTR)* were conducted in accordance with the Animal Welfare Act and with permission of the Landesdirektion Sachsen, Germany (DD24-5131/346/11, DD24-5131/346/12, DD24.1-5131/476/2, TVV21/2018 and all corresponding amendments). Zebrafish husbandry and experiments with all other transgenic lines was performed under standard conditions in accordance with institutional (Université Libre de Bruxelles (ULB)) and national ethical and animal welfare guidelines and regulation, which were approved by the ethical committee for animal welfare (CEBEA) from the Université Libre de Bruxelles (protocols 578N-579N).

### Dissection of the zebrafish thyroid gland

The dissection of thyroid gland in zebrafish was performed by using the ventral aorta as a reference (Fig. 1A, B). In zebrafish, the thyroid follicles sit loosely in soft tissue around the ventral aorta. Ventral aorta connects to the outflow tract that further joins with the heart ventricle. During dissociation, cells connected to the ventral aorta, including parts of zebrafish gills (Fig. 1C) were kept intact to avoid injuring the organ during dissociation.

In detail, zebrafish were euthanized in 0.2% Tricaine (MS-222, Sigma E10521) solution. Using fine forceps, the lower jaw was separated from the upper jaw and disconnected from the gut by pinching near the gills. The dissected tissue was carefully cleaned by removing muscle, skin, pectoral fin and lateral cartilages of the lower jaw. The cleaned tissue when observed from the ventral side under brightfield clearly shows the ventral aorta as a thick pink blood vessel extending from the heart towards the basibranchial cartilage (Fig. 1A). Next, the surrounding gills are pinched off using fine forceps, taking care to keep the ventral aorta intact (Fig. 1B). This leaves small parts of gills connected to the ventral aorta (Fig. 1C). Lastly, the ventral aorta is disconnected from the outflow tract by pinching with fine forceps (dashed line in Fig. 1B).

### Single cell suspension of zebrafish thyroid gland

Single cell suspension of zebrafish thyroid gland was performed by adapting the cell dissociation protocol outlined in Singh et al., Scientific Reports, 2018 (46). In brief, the thyroid gland was collected and dissociated into single cells by incubation in TrypLE (ThermoFisher, 12563029) with 0.1% Pluronic F-68 (ThermoFisher, 24040032) at 37 °C in a benchtop shaker set at 450 rpm for 45 min. Following dissociation, TrypLE was inactivated with 10% FBS, and the cells pelleted by centrifugation at 500g for 10min at 4 °C. The supernatant was carefully discarded and the pellet re-suspended in 500 uL of HBSS (without Ca, Mg) + 0.1% Pluronic F-68. To remove debris, the solution was passed over a 30 μm cell filter (Miltenyi Biotec, 130-041-407). To remove dead cells, calcein violet (ThermoFisher, C34858) was added at a final concentration of 1 μM and the cell suspension incubated at room temperature for 20 minutes. The single cell preparation was sorted with the appropriate gates, including excitation with UV (405 nm) laser for identification of alive cells (calcein+) (Fig. 1F – G). FACS was performed through 100 μm nozzle.

### Single-cell profiling of the zebrafish thyoid gland

For single-cell RNA-seq of the zebrafish thyroid gland using the 10X Genomics platform, cell suspension was prepared as mentioned above from the thyroid glands of six 2 month post fertilization and six 8 month post-fertilization *Tg(tg:nls-mVenus-T2A-NTR)* animals. The cell suspension was adjusted with Hanks’ Balanced Salt Solution (without calcium and magnesium) to a density of 800cells/μl, and diluted with nuclease-free water according to the manufacturer’s instructions to yield 12,000 cells. Subsequently, the cells were carefully mixed with reverse transcription mix before loading the cells on the 10X Genomics Chromium system (10). After the gel emulsion bead suspension underwent the reverse transcription reaction, emulsion was broken and DNA purified using Silane beads. The complementary DNA was amplified with 10 cycles, following the guidelines of the 10x Genomics user manual. The 10X Genomics single cell RNA-seq library preparation—involving fragmentation, dA tailing, adapter ligation and indexing PCR—was performed based on the manufacturer’s protocol. After quantification, the libraries were sequenced on an Illumina NextSeq 550 machine using a HighOutput flowcell in paired-end mode (R1: 26 cycles; I1: 8 cycles; R2: 57 cycles), thus generating ~45mio fragments. The raw sequencing data were then processed with the ‘count’ command of the Cell Ranger software (v.2.1.0) provided by 10x Genomics with the option ‘–expect-cells’ set to 10,000 (all other options were used as per default).

To build the reference for Cell Ranger, zebrafish genome (GRCz10) as well as gene annotation (Ensembl 87) were downloaded from Ensembl and the annotation was filtered with the ‘mkgtf’ command of Cell Ranger (options: ‘– attribute=gene_biotype:protein_coding– attribute=gene_biotype:lincRNA – attribute=gene_biotype:antisense’). Genome sequence and filtered annotation were then used as input to the ‘mkref’ command of Cell Ranger to build the appropriate Cell Ranger Reference.

### Analysis of single-cell RNA-Seq. of the zebrafish thyroid gland

The raw data generated from 10x Chromium pipeline was clustered using Seurat 2.3.4 (47) using the recommended analysis pipeline. Briefly, the raw data as UMI-counts was log-normalized, regressed to remove the effect of library size and mitochondrial counts, and scaled. Highly variable genes were identified for PCA analysis and graph-based clustering using shared nearest neighbour (SNN). For clustering, the first five principal components (PCs) were utilized as they displayed significant deviation from uniform distribution as accessed by JackStraw analysis. Further, a resolution of 0.3 for SNN was used for clustering. Marker genes identified for each cluster were used to classify the cell-type. The thyrocyte cluster was isolated and sub-clustered using the first three PCs and 0.3 resolution to identify and label sub-populations.

### Development of intercellular signaling network

Development of intercellular signaling network for zebrafish was performed as described in Cosacak et al. (48). Briefly, ligands expressed in 20% of a cell-population were identified. A connection between cell-type and thyrocyte was made if the expression of the corresponding receptor was identified in 20% of thyrocytes. The connectome contains secreted and membrane-tethered ligands. For cell-types that do not physically contact the thyrocytes (gills, NFE and stroma), membrane-tethered ligands were manually removed from the connectome.

### Background correction for thyrocyte gene expression

Supervised background correction for the thyrocyte population was performed using DecontX (49). As input, normalized data and clustering information from Seurat was used. The method using Bayesian approach to model gene expression as a mixture of expression in the expected cell-population plus background expression accessed from remaining cell-types. Background noise is removed, which likely resembles free mRNA released from injured and ruptured cells. As background correction required clustering information, the corrected data was not utilized for re-clustering to avoid circular use of the data.

### Genetic entropy analysis for thyrocyte population

Quantification of genetic entropy was performed using ROGUE (Ratio of Global Unshifted Entropy) (50). As input, raw counts of thyrocytes that passed quality control were used. Default parameters were used for analysis. The algorithm provided a measure of entropy (degree of uncertainty / heterogeneity), along with p-value, within the population.

### Data Availability

The raw 10X data, along with tabulated count data are available publicly from GEO under accession number GSE133466. The atlas for online browsing is available at https://sumeet.shinyapps.io/zfthyroid/.

### Generation of knock-in *pax2a*^*pax2a-T2A-mKO2*^ zebrafish line

For generation of *pax2a* reporter line, we designed a single-guide RNA (sgRNA) targeting the STOP codon of the *pax2a* coding sequence (GCTGCGATGGTAACTAGTGG). We then generated a donor construct in which the sequence encoding for the monomeric Kusabira orange (mKO2) protein was fused to a viral 2A peptide linker. This reporter cassette was flanked by left (1000bp) and right (2000bp) homology arms of the *pax2a* genomic DNA region around the stop codon therefore preventing the sgRNA from cutting the donor construct. sgRNA design, production and validation were done as previously described (27, 51). Wild-type embryos were injected with 3 nL of the injection mix containing the sgRNA (final concentration 80 ng/μL), the donor construct (final concentration 7.5 ng/μL), the protein Cas9 (recombinant cas protein from *S. pyogenes* PNA Bio CP01, final concentration 100 ng/μL) and KCL (final concentration 200 mM). Upon homologous recombination of this reporter construct in the endogenous locus, *pax2a*-expressing cells were fluorescently labelled by mKO2. This *pax2a*^*pax2a-T2A-mKO2*^ line is referenced as *pax2a*^mKO2^ in the text.

### Generation of *pax2a* crispants

Somatic mutagenesis of pax2a gene was carried out exactly as mentioned in Trubiroha et al., Scientific Reports, 2018 (27). Briefly, sgRNA targeting the exon 2 of *pax2a* was generated as described in the publication. Following the strategy described in the publication, Cas9 protein along with sgRNA was injected in one-cell stage of zebrafish embryos for disruption of *pax2a* gene. Non-injected animals were used as controls.

### Tissue collection

To facilitate confocal imaging of the thyroid gland, the organ was manually dissected from fish as previously described and fixed. Fish were killed in Tricaine followed by dissection of the gland, which was fixed by immersion in 4% paraformaldehyde (PFA) + 1% Triton-X overnight at 4 °C. The gland was washed 2 – 3 times in PBS to remove PFA before proceeding.

### Quantification of proportion of thyrocytes within the dissected tissue

To quantify the proportion of thyrocytes within the dissected tissue, the gland was dissected from *Tg(tg:nls-EGFP)* animals and fixed as described above. The fixed tissue was permeabilized by three washes with 1% PBT (1x PBS + 1 % Triton-X-100). Nuclei were stained by immersing the tissue in 1 μg / ml Hoechst prepared in 1x PBS for two hours at room temperature. The tissue was immersed in 30% sucrose solution overnight at 4 °C, embedded in Tissue Freezing Medium (Leica 14020108926) and frozen at −80 °C. Thin sections (8 μm) were obtained using cryostat (Leica CM3050 S), collected on frosted glass slides (Thermo Scientific 12362098) and covered with glass coverslip of #1 thickness (Carl Roth GmbH NK79.1) using mounting media (Dako S3023). The sections were imaged on Zeiss LSM 780 confocal microscope. Confocal images were analyzed in Fiji using the following step: threshold using ‘IsoData’ to distinguish signal from background, ‘watershed’ transformation to separate joined nuclei and ‘measure’ function to obtain nuclei count. With this, the green channel (number of thyrocyte nuclei) and blue channel (total number of nuclei) was measured for seven transverse sections obtained from three animals. Percentage was calculated by taking the ratio of thyrocyte nuclei to total nuclei.

### Immunofluorescence and image acquisition

Whole-mount immunofluorescence was performed on thyroid gland collected as described above. The collected samples were permeabilized in 1% PBT (Triton-X-100) and blocked in 4% PBTB (BSA). Primary and secondary antibody stainings were performed overnight at 4 °C. Primary antibodies used in this study were anti-PAX2A (rabbit, Genetex GTX128127) at 1:250, anti-EGFP (chicken, Abcam ab13970) at 1:1000, anti-E-Cadherin (mouse, BD bioscience cat 610181) at 1:200, anti-monomeric Kusabira-Orange 2 (mouse, MBL amalgaam M-168-3M) at 1:200, anti-monomeric Kusabira-Orange 2 (rabbit, MBL amalgam PM051M) at 1:250 and anti-p63 (mouse, Santa Cruz Biotechnology 4A4) at 1:200. Secondary antibodies at 1:250 dilutions used in this study were Alexa Fluor 488 anti-chicken (Jackson ImmunoResearch laboratories 703-545-155), Alexa Fluor 647 anti-rabbit (Jackson ImmunoResearch laboratories 711-605-152), Alexa Fluor 647 anti-mouse (Jackson ImmunoResearch laboratories 715-605-150), Cy™3-conjugated anti-rabbit (Jackson ImmunoResearch laboratories 711-165-152) and Cy™3-conjugated anti-mouse (Jackson ImmunoResearch laboratories 715-165-150). When needed nuclei were staining using DAPI at a 1:1000 dilution. Samples were mounted in NuSieve^TM^ GTG^TM^ Agarose (Lonza cat50080) and imaged on a glass bottom FluoroDish^TM^ (WPI FD3510-100) using a Zeiss LSM 780 confocal microscope or Leica DMI 6000b microscope. ImageJ was used to add scale bars and PowerPoint was used for adding arrows and labels.

### FACS-based reporter analysis

For analysing the levels of *pax2a*^*mKO2*^ by FACS, single-cell suspension from the thyroid gland of 5 mpf *Tg(tg:nls-GFP)*; *pax2a*^*mKO2*^ animals was prepared as described earlier and stained with 1 μM calcein violet (ThermoFisher, C34858). Cells were sorted and analyzed using FACS-Aria II (BD Bioscience). Thyrocytes were selected by gating for calcein+ GFP+ population, and mKO2 expression level recorded for analysis.

### Gene Ontology (GO) Analysis

Gene ontology (GO) analysis was performed using DAVID (52). The list of genes was uploaded on the web browser of DAVID and statistically significant (p-value < 0.05) GO terms were identified using default parameters.

### Statistical analysis

Statistical analysis was performed using R. No animals were excluded from analysis. Blinding was not performed during analysis. Analysis of normal distribution was not performed.

## Acknowledgements

We thank members of the Costagliola and Singh lab for comments on the manuscript, members of Center for Regenerative Therapies Dresden (CRTD) fish, FACS and sequencing facility, and members of IRIBHM fish facility for technical assistance. We thank J.-M. Vanderwinden from the Light Microscopy Facility and Christine Dubois from the FACS facility for technical assistance at ULB. We are grateful to Priyanka Oberoi for illustrations. P.G is Fund for Research in the Industry and the Agriculture (FRIA) Research fellow; M.S. is FNRS Research Fellow (34985615 - THYSCEFA); S.C. is FNRS Senior Research Associate. V.D. acknowledges grants from the Fond Naets (J1813300), the Fondation Contre le Cancer (2016-093) and FNRS (EQP/OL U.N019.19, J006120F). Work by M.B., C.L. and G.K. was supported by grants to M.B. from the Deutsche Forschungsgemeinschaft and European Union (European Research Council AdG Zf-BrainReg). Work by S.P.S. was supported by MISU funding from the FNRS (34772792 – SCHISM). This work was supported by grants from the Belgian National Fund for Scientific Research (FNRS) (FRSM 3-4598-12; CDR-J.0145.16, GEQ U.G030.19), the Fonds d’Encouragement à la Recherche de l’Université Libre de Bruxelles (FER-ULB).

## Author contribution

S.P.S. conceptualized the project. N.N., G.K., C.L. and M.B. provided reagents and animals for single-cell RNA-Sequencing. S.P.S., S.R., A.K., J.B., and A.P. performed the single-cell RNA-Sequencing. S.P.S., S.E.E., V.D., S.C. analysed and interpreted the data. S.P.S. developed the online browser. P.G. and B.H. generated the *pax2a* knock-in line. P.G. and M.S. analysed the *pax2a* reporter line, M.P.M. and I.G.S. collected immunofluorescence images. S.P.S. wrote the first draft and S.C., P.G., S.E.E edited the manuscript. S.P.S. and S.C. acquired funding for the project. All authors read and approved the final manuscript.

## Conflict of interest

The authors declare that they have no conflict of interest.

## Notes

#### Summary of Updates

Manuscript revised in response to peer-reviews received from Review Commons. In this revision, dissection of the thyroid gland from zebrafish has been clarified, cell types adjacent to thyroid follicles have been characterised using immunofluorescence and transcriptional heterogeneity in pax2a expression in thyroid follicular cells verified using antibody staining. For this, Figures, Results and Methods Sections have been updated. In addition, Authors who contributed to the revision have been added.

https://sumeet.shinyapps.io/zfthyroid/

## References

1. Taylor PN, Albrecht D, Scholz A, Gutierrez-Buey G, Lazarus JH, Dayan CM, Okosieme OE 2018 Global epidemiology of hyperthyroidism and hypothyroidism. Nat Rev Endocrinol 14:301–316.

2. Brix K, Lemansky P, Herzog V 1996 Evidence for extracellularly acting cathepsins mediating thyroid hormone liberation in thyroid epithelial cells. Endocrinology 137:1963–74.

3. Alt B, Reibe S, Feitosa NM, Elsalini OA, Wendl T, Rohr KB 2006 Analysis of origin and growth of the thyroid gland in zebrafish. Dev Dyn 235:1872–1883.

4. Linehan SA, Martínez-Pomares L, da Silva RP, Gordon S 2001 Endogenous ligands of carbohydrate recognition domains of the mannose receptor in murine macrophages, endothelial cells and secretory cells; potential relevance to inflammation and immunity. Eur J Immunol 31:1857–66.

5. Nonidez JF 1931 Innervation of the thyroid gland. II. Origin and course of the thyroid nerves in the dog. Am J Anat 48:299–329.

6. Svensson V, Vento-Tormo R, Teichmann SA 2018 Exponential scaling of single-cell RNA-seq in the past decade. Nat Protoc 13:599–604.

7. McMenamin SK, Bain EJ, McCann AE, Patterson LB, Eom DS, Waller ZP, Hamill JC, Kuhlman JA, Eisen JS, Parichy DM 2014 Thyroid hormone-dependent adult pigment cell lineage and pattern in zebrafish. Science (80-) 345:1358–1361.

8. AlJanahi AA, Danielsen M, Dunbar CE 2018 An Introduction to the Analysis of Single-Cell RNA-Sequencing Data. Mol Ther Methods Clin Dev 10:189–196.

9. Macosko EZ, Basu A, Satija R, Nemesh J, Shekhar K, Goldman M, Tirosh I, Bialas AR, Kamitaki N, Martersteck EM, Trombetta JJ, Weitz DA, Sanes JR, Shalek AK, Regev A, McCarroll SA 2015 Highly Parallel Genome-wide Expression Profiling of Individual Cells Using Nanoliter Droplets. Cell 161:1202–1214.

10. Zheng GXY, Terry JM, Belgrader P, Ryvkin P, Bent ZW, Wilson R, Ziraldo SB, Wheeler TD, McDermott GP, Zhu J, Gregory MT, Shuga J, Montesclaros L, Underwood JG, Masquelier DA, Nishimura SY, Schnall-Levin M, Wyatt PW, Hindson CM, Bharadwaj R, Wong A, Ness KD, Beppu LW, Deeg HJ, McFarland C, Loeb KR, Valente WJ, Ericson NG, Stevens EA, Radich JP, Mikkelsen TS, Hindson BJ, Bielas JH 2017 Massively parallel digital transcriptional profiling of single cells. Nat Commun 8:14049.

11. Kivioja T, Vähärautio A, Karlsson K, Bonke M, Enge M, Linnarsson S, Taipale J 2011 Counting absolute numbers of molecules using unique molecular identifiers. Nat Methods 9:72–4.

12. Islam S, Zeisel A, Joost S, La Manno G, Zajac P, Kasper M, Lönnerberg P, Linnarsson S 2014 Quantitative single-cell RNA-seq with unique molecular identifiers. Nat Methods 11:163–6.

13. Maaten L van der, Hinton G 2008 Visualizing data using t-SNE. J Mach Learn Res 9:2579–2605.

14. Sánchez-Iranzo H, Galardi-Castilla M, Sanz-Morejón A, González-Rosa JM, Costa R, Ernst A, Sainz de Aja J, Langa X, Mercader N 2018 Transient fibrosis resolves via fibroblast inactivation in the regenerating zebrafish heart. Proc Natl Acad Sci U S A 115:4188–4193.

15. Denton CP, Zheng B, Shiwen X, Zhang Z, Bou-Gharios G, Eberspaecher H, Black CM, de Crombrugghe B 2001 Activation of a fibroblast-specific enhancer of the proalpha2(I) collagen gene in tight-skin mice. Arthritis Rheum 44:712–22.

16. Lisse TS, Middleton LJ, Pellegrini AD, Martin PB, Spaulding EL, Lopes O, Brochu EA, Carter E V, Waldron A, Rieger S 2016 Paclitaxel-induced epithelial damage and ectopic MMP-13 expression promotes neurotoxicity in zebrafish. Proc Natl Acad Sci U S A 113:E2189–98.

17. Reischauer S, Levesque MP, Nüsslein-Volhard C, Sonawane M 2009 Lgl2 executes its function as a tumor suppressor by regulating ErbB signaling in the zebrafish epidermis. PLoS Genet 5:e1000720.

18. Barbieri CE, Pietenpol JA 2006 p63 and epithelial biology. Exp Cell Res 312:695–706.

19. Ramilowski JA, Goldberg T, Harshbarger J, Kloppman E, Lizio M, Satagopam VP, Itoh M, Kawaji H, Carninci P, Rost B, Forrest ARR 2015 A draft network of ligand-receptor-mediated multicellular signalling in human. Nat Commun 6:7866.

20. Brix K, Linke M, Tepel C, Herzog V 2001 Cysteine proteinases mediate extracellular prohormone processing in the thyroid. Biol Chem 382:717–25.

21. Linke M 2002 Trafficking of lysosomal cathepsin B--green fluorescent protein to the surface of thyroid epithelial cells involves the endosomal/lysosomal compartment. J Cell Sci 115:4877–4889.

22. Wendl T, Lun K, Mione M, Favor J, Brand M, Wilson SW, Rohr KB 2002 Pax2.1 is required for the development of thyroid follicles in zebrafish. Development 129:3751–60.

23. Porazzi P, Calebiro D, Benato F, Tiso N, Persani L 2009 Thyroid gland development and function in the zebrafish model. Mol Cell Endocrinol 312:14–23.

24. Pfeffer PL, Gerster T, Lun K, Brand M, Busslinger M 1998 Characterization of three novel members of the zebrafish Pax2/5/8 family: dependency of Pax5 and Pax8 expression on the Pax2.1 (noi) function. Development 125:3063–74.

25. Mansouri A, Chowdhury K, Gruss P 1998 Follicular cells of the thyroid gland require Pax8 gene function. Nat Genet 19:87–90.

26. Kesavan G, Chekuru A, Machate A, Brand M 2017 CRISPR/Cas9-Mediated Zebrafish Knock-in as a Novel Strategy to Study Midbrain-Hindbrain Boundary Development. Front Neuroanat 11:52.

27. Trubiroha A, Gillotay P, Giusti N, Gacquer D, Libert F, Lefort A, Haerlingen B, De Deken X, Opitz R, Costagliola S 2018 A Rapid CRISPR/Cas-based Mutagenesis Assay in Zebrafish for Identification of Genes Involved in Thyroid Morphogenesis and Function. Sci Rep 8:5647.

28. Liang S, Johansson E, Barila G, Altschuler DL, Fagman H, Nilsson M 2018 A branching morphogenesis program governs embryonic growth of the thyroid gland. Development 145:dev146829.

29. Guzman A, Ramos-Balderas JL, Carrillo-Rosas S, Maldonado E 2013 A stem cell proliferation burst forms new layers of P63 expressing suprabasal cells during zebrafish postembryonic epidermal development. Biol Open 2:1179–1186.

30. Yang A, Schweitzer R, Sun D, Kaghad M, Walker N, Bronson RT, Tabin C, Sharpe A, Caput D, Crum C, McKeon F 1999 p63 is essential for regenerative proliferation in limb, craniofacial and epithelial development. Nature 398:714–8.

31. Baber EC 1876 XXI. Contributions to the minute anatomy of the thyroid gland of the dog. Philos Trans R Soc London 166:557–568.

32. Getzowa S 1907 Über die Glandula parathyreodeaa, intrathyreoideale Zellhaufen derselben und Reste des postbranchialen Körpers. Virchows Arch Pathol Anat Physiol Klin Med 188:181–235.

33. Harach HR 1988 Solid cell nests of the thyroid. J Pathol 155:191–200.

34. Ríos Moreno MJ, Galera-Ruiz H, De Miguel M, López MIC, Illanes M, Galera-Davidson H 2011 Inmunohistochemical profile of solid cell nest of thyroid gland. Endocr Pathol 22:35–9.

35. Rasmussen JP, Sack GS, Martin SM, Sagasti A 2015 Vertebrate epidermal cells are broad-specificity phagocytes that clear sensory axon debris. J Neurosci 35:559–70.

36. Neill T, Schaefer L, Iozzo R V 2012 Decorin: a guardian from the matrix. Am J Pathol 181:380–7.

37. Martínez-Aguilar J, Clifton-Bligh R, Molloy MP 2016 Proteomics of thyroid tumours provides new insights into their molecular composition and changes associated with malignancy. Sci Rep 6:23660.

38. Planck T, Shahida B, Sjögren M, Groop L, Hallengren B, Lantz M 2014 Association of BTG2, CYR61, ZFP36, and SCD gene polymorphisms with Graves’ disease and ophthalmopathy. Thyroid 24:1156–61.

39. Marinò M, McCluskey RT 2000 Megalin-mediated transcytosis of thyroglobulin by thyroid cells is a calmodulin-dependent process. Thyroid 10:461–9.

40. Tu D-G, Chang W-W, Jan M-S, Tu C-W, Lu Y-C, Tai C-K 2016 Promotion of metastasis of thyroid cancer cells via NRP-2-mediated induction. Oncol Lett 12:4224–4230.

41. Antonica F, Kasprzyk DF, Opitz R, Iacovino M, Liao X-H, Dumitrescu AM, Refetoff S, Peremans K, Manto M, Kyba M, Costagliola S 2012 Generation of functional thyroid from embryonic stem cells. Nature 491:66–71.

42. Chopra K, Ishibashi S, Amaya E 2019 Zebrafish duox mutations provide a model for human congenital hypothyroidism. Biol Open 8.

43. Anelli V, Villefranc JA, Chhangawala S, Martinez-McFaline R, Riva E, Nguyen A, Verma A, Bareja R, Chen Z, Scognamiglio T, Elemento O, Houvras Y 2017 Oncogenic BRAF disrupts thyroid morphogenesis and function via twist expression. Elife 6.

44. Choi J, Dong L, Ahn J, Dao D, Hammerschmidt M, Chen J-N 2007 FoxH1 negatively modulates flk1 gene expression and vascular formation in zebrafish. Dev Biol 304:735–744.

45. Ellett F, Pase L, Hayman JW, Andrianopoulos A, Lieschke GJ 2011 mpeg1 promoter transgenes direct macrophage-lineage expression in zebrafish. Blood 117:e49–56.

46. Singh SP, Janjuha S, Chaudhuri S, Reinhardt S, Kränkel A, Dietz S, Eugster A, Bilgin H, Korkmaz S, Zararsız G, Ninov N, Reid JE 2018 Machine learning based classification of cells into chronological stages using single-cell transcriptomics. Sci Rep 8:17156.

47. Butler A, Hoffman P, Smibert P, Papalexi E, Satija R 2018 Integrating single-cell transcriptomic data across different conditions, technologies, and species. Nat Biotechnol doi: 10.1101/164889.

48. Cosacak MI, Bhattarai P, Reinhardt S, Petzold A, Dahl A, Zhang Y, Kizil C 2019 Single-Cell Transcriptomics Analyses of Neural Stem Cell Heterogeneity and Contextual Plasticity in a Zebrafish Brain Model of Amyloid Toxicity. Cell Rep 27:1307–1318.e3.

49. Yang S, Corbett SE, Koga Y, Wang Z, Johnson WE, Yajima M, Campbell JD 2020 Decontamination of ambient RNA in single-cell RNA-seq with DecontX. Genome Biol 21:57.

50. Liu B, Li C, Li Z, Ren X, Zhang Z 2019 ROGUE: an entropy-based universal metric for assessing the purity of single cell population. bioRxiv 819581.

51. Varshney GK, Pei W, LaFave MC, Idol J, Xu L, Gallardo V, Carrington B, Bishop K, Jones M, Li M, Harper U, Huang SC, Prakash A, Chen W, Sood R, Ledin J, Burgess SM 2015 High-throughput gene targeting and phenotyping in zebrafish using CRISPR/Cas9. Genome Res 25:1030–42.

52. Huang D, Sherman BT, Tan Q, Collins JR, Alvord WG, Roayaei J, Stephens R, Baseler MW, Lane HC, Lempicki RA 2007 The DAVID Gene Functional Classification Tool: a novel biological module-centric algorithm to functionally analyze large gene lists. Genome Biol 8:R183.

