## Supplementary Data for "Single-cell transcriptome analysis reveals cell-cell communication and thyrocyte diversity in the zebrafish thyroid gland"

**Includes**

Supplementary Figures 1 – 8

Supplementary Tables 1 – 4 Legend

Supplementary Movie 1 Legend

### Supplementary Figures

#### Supplementary Figure 1

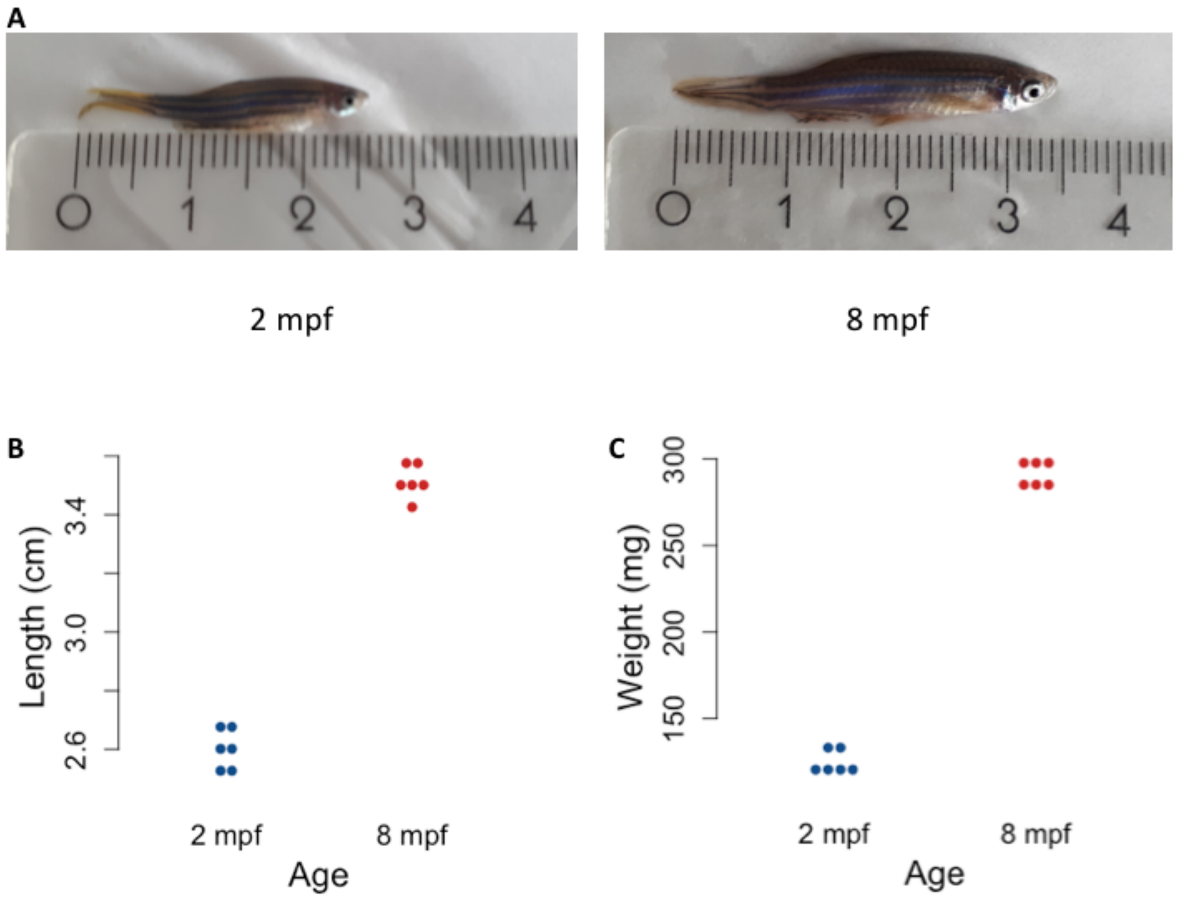


**Supplementary Figure 1: Physical characteristics of zebrafish at 2 mpf and 8 mpf.** **(A)** Representative images of zebrafish at 2 mpf (left) and 8 mpf (right). **(B)** Dotplot representing the length of individual animals at each stage. Y-axis represents the length of six zebrafish from mouth to end of fin in cm. **(C)** Dotplot representing the weight of individual animals at each stage. Y-axis represents the weight of six zebrafish in mg.

**Supplementary Figure 2**

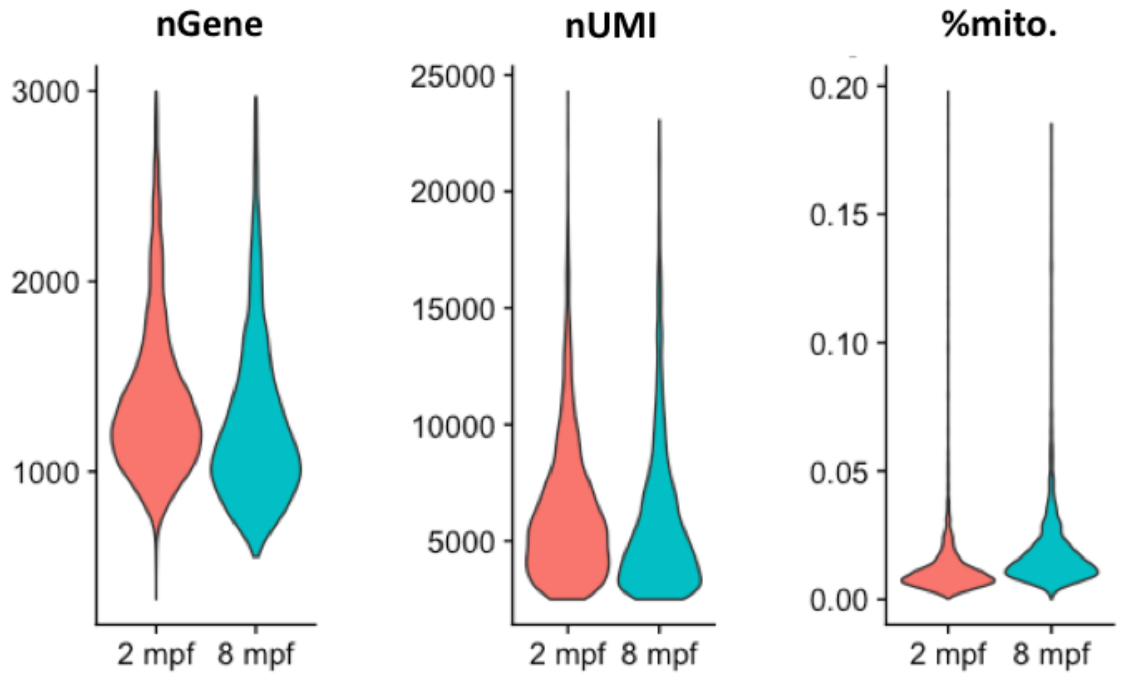


**Supplementary Figure 2: Quality control parameters for cells present in the zebrafish thyroid gland atlas**

Violin plots depicting the number of genes (nGene), number of unique RNA molecules detected (nUMI – number of Unique Molecular Identifier) and the percentage of reads mapped onto the mitochondrial genome for the cells profiled in the zebrafish thyroid gland atlas.

**Supplementary Figure 3**

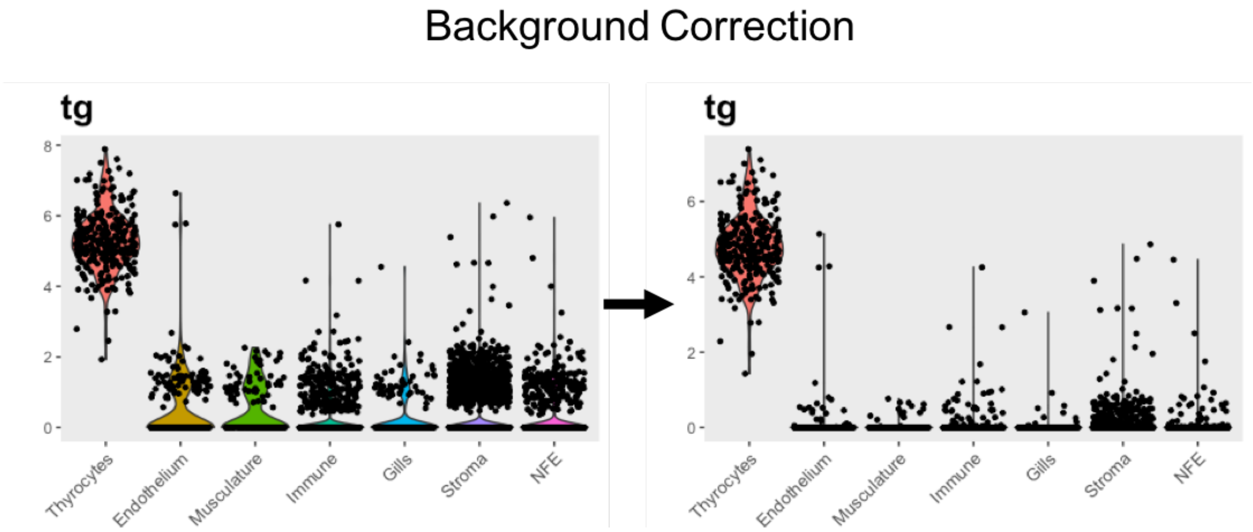


**Supplementary Figure 3: Background correction reduces expression noise**

Single-cell RNA-Seq. suffers from background noise due to cross-contamination from free mRNA released from ruptured and injured cell. Noise reduction lead to reduced *tg* expression in non-thyrocyte cell-populations.

**Supplementary Figure 4**

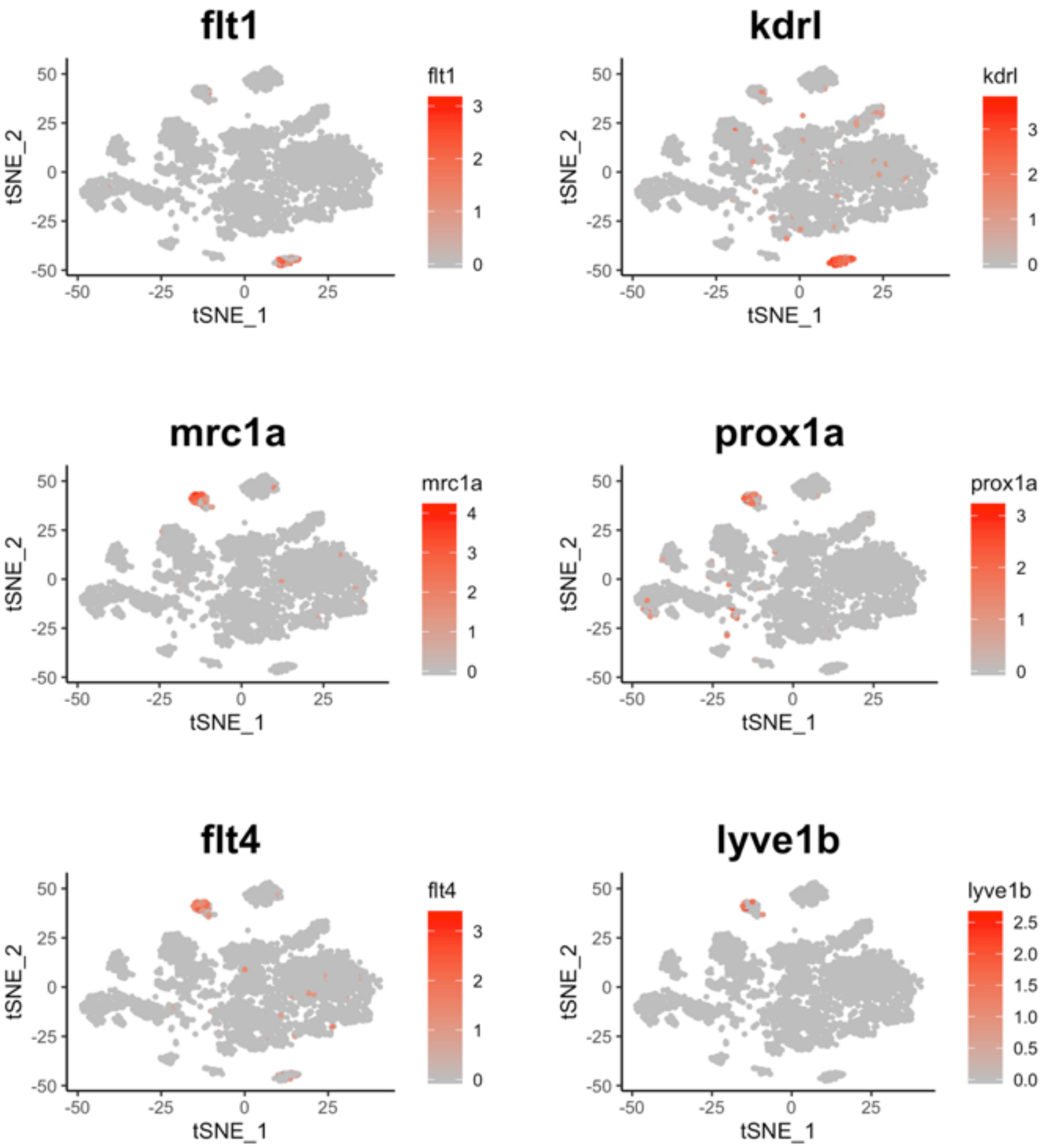


**Supplementary Figure 4: Cells belonging to blood vessels and lymphatic vessels are present in the zebrafish thyroid gland atlas**

T-SNE plot overlaid with gene expression for genes specific to blood vessels (*flt1* and *kdrl*) and lymphatic vessles (*mrc1a*, *prox1a*, *flt4* and *lyve1b*).

**Supplementary Figure 5**

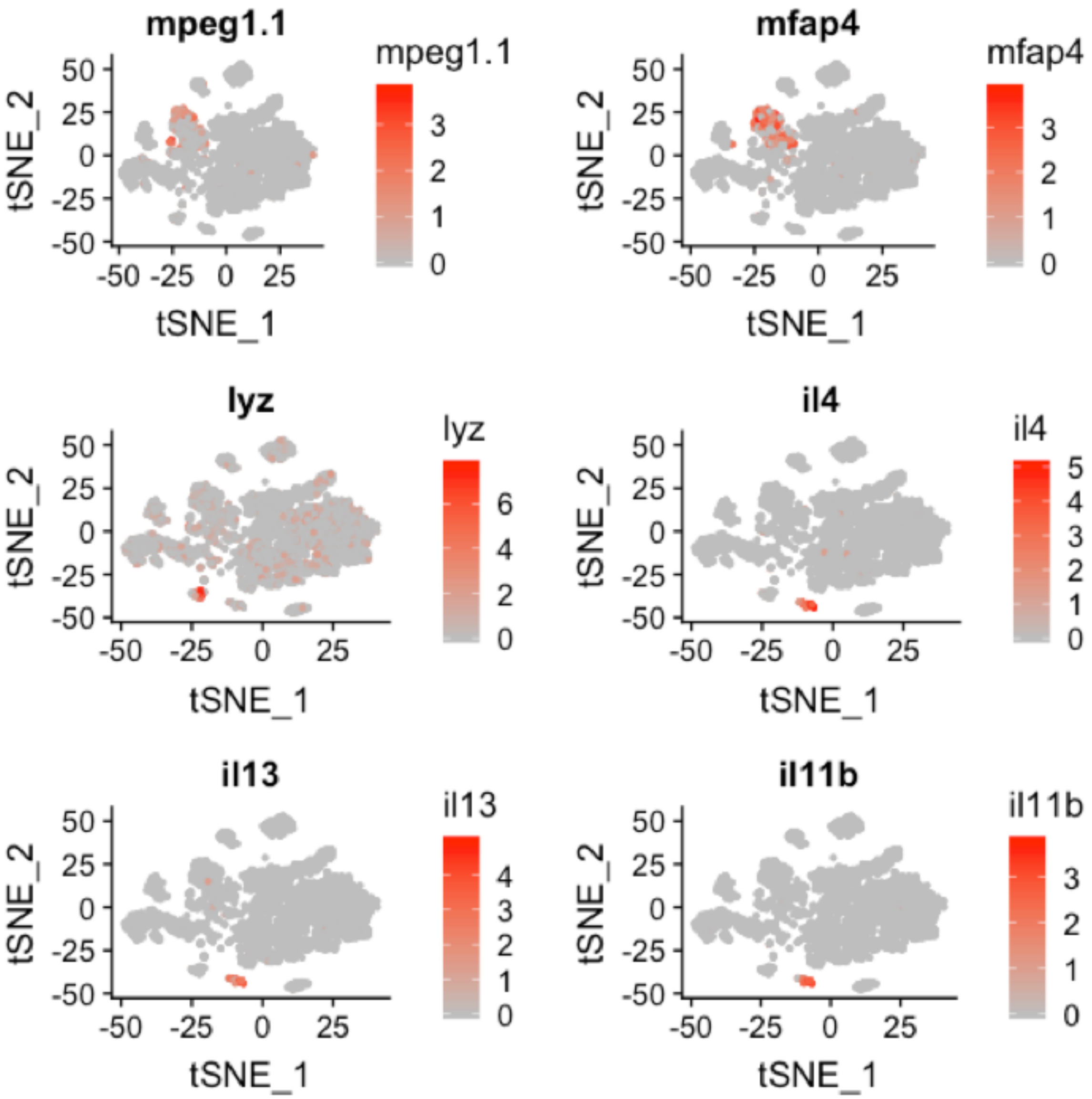


**Supplementary Figure 5: Cells belonging to diverse hematopoietic lineage are present in the zebrafish thyroid gland atlas**

T-SNE plot overlaid with gene expression for genes specific to macrophages (*mpeg1.1* and *mfap4*), neutrophils (*lyz*) and lymphocytes (*il4*, *il13* and *il11b*).

**Supplementary Figure 6**

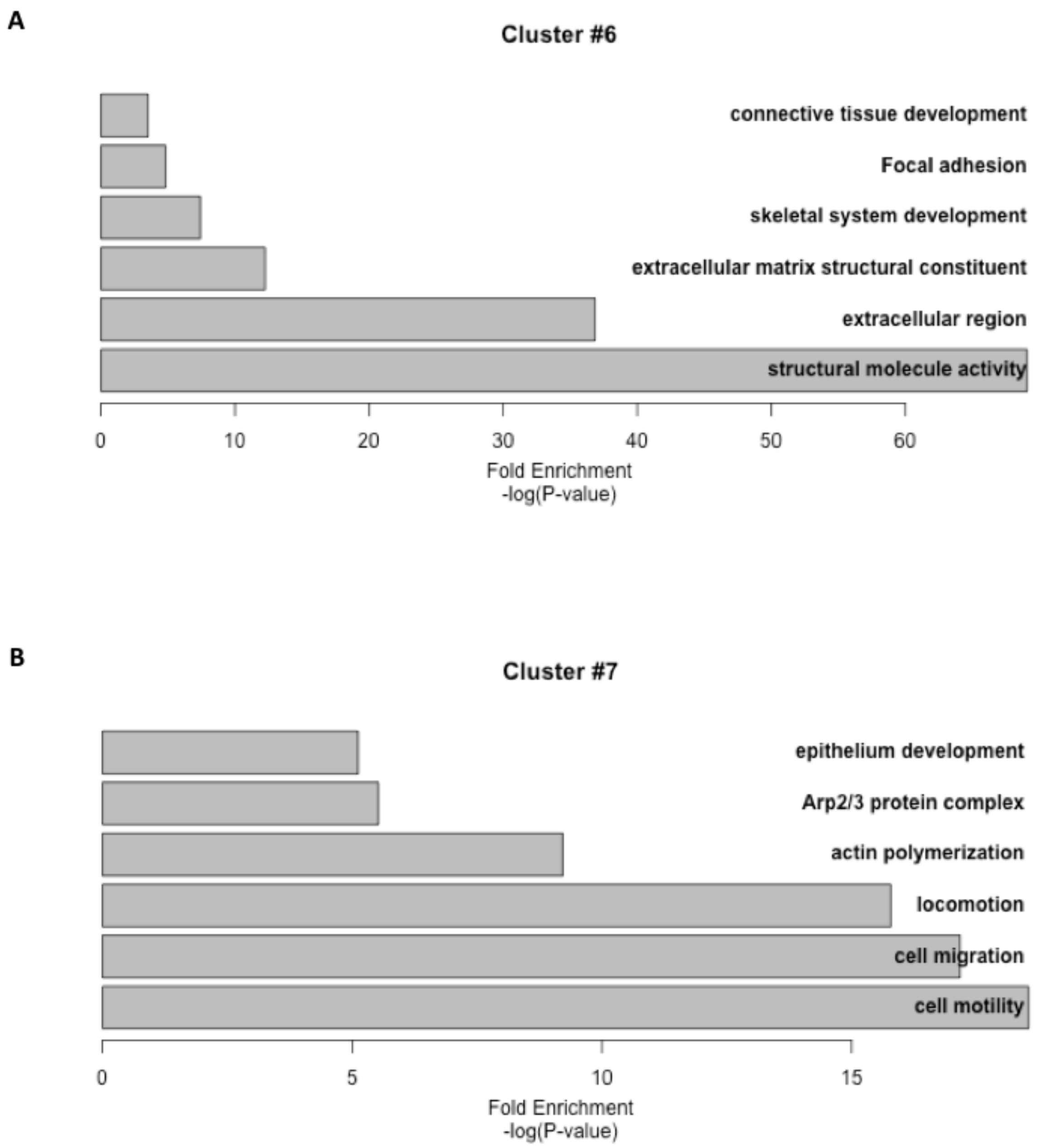


**Supplementary Figure 6: GO analysis for Cluster #6 and #7 marker genes**

Barplot depicting the GO categories identified for marker genes enriched in Cluster #6 **(A)** and Cluster #7**(B)**. X-axis represents a negative log of p-value.

**Supplementary Figure 7**

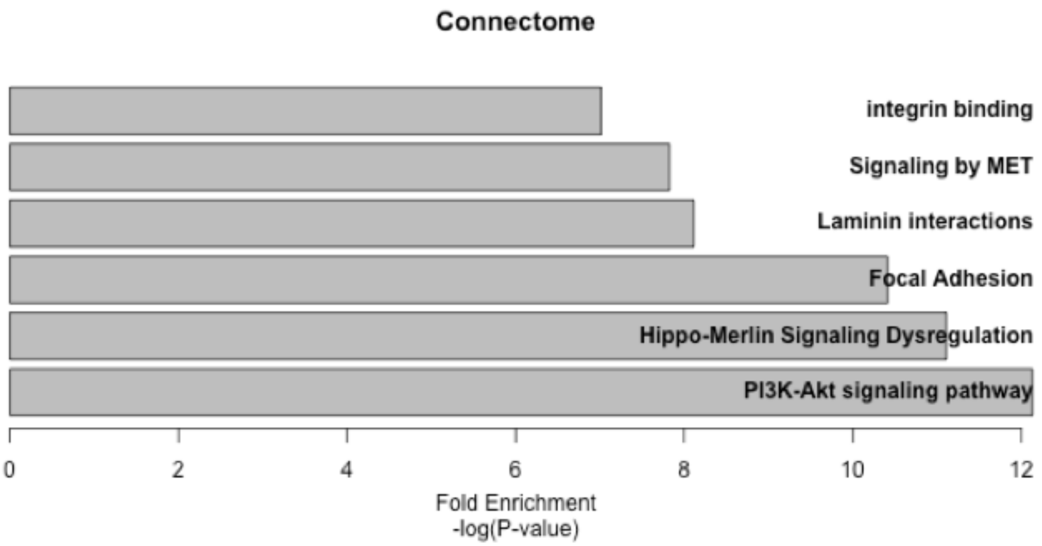


**Supplementary Figure 7: GO analysis for zebrafish thyroid gland connectome**

Barplot depicting the GO categories for ligands expressed in different cell-types present in the thyroid gland atlas and their respective receptors in thyrocytes. X-axis represents a negative log of p-value.

### Supplementary Figure 8

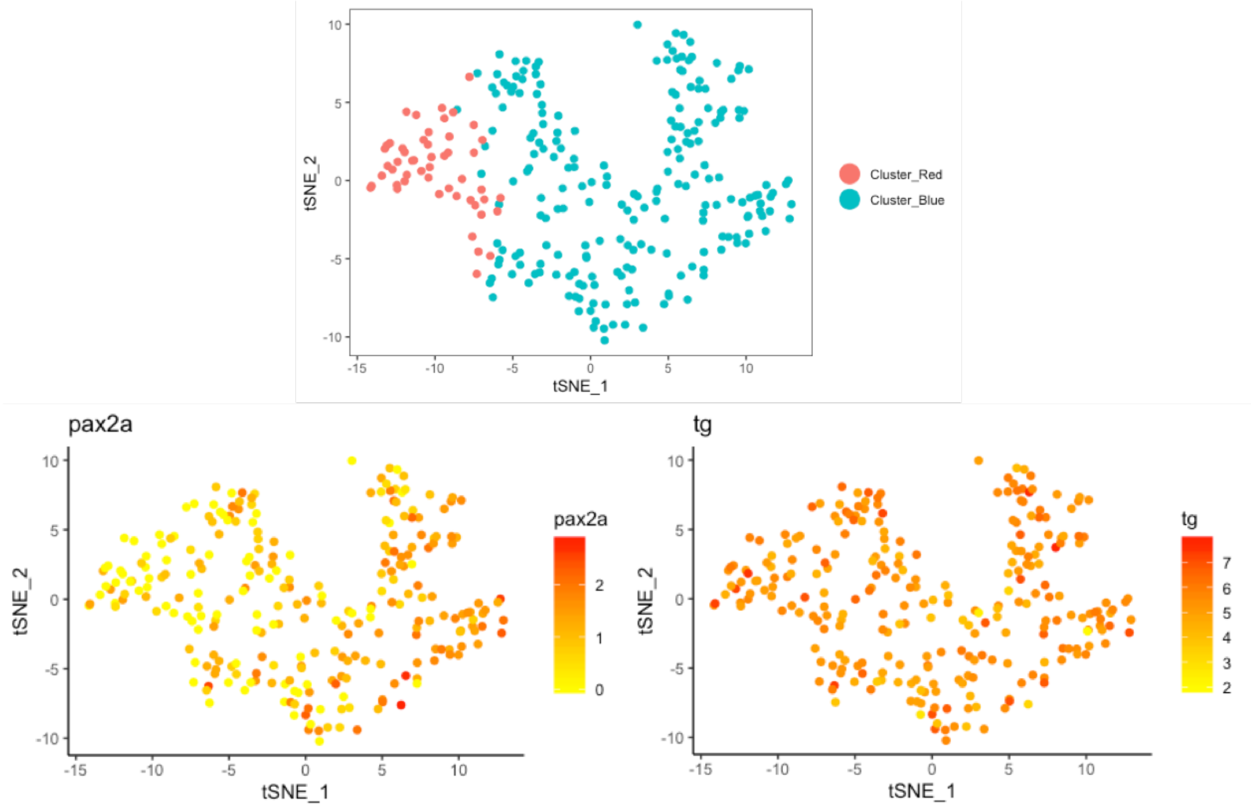


**Supplementary Figure 8: Expression of *pax2a* and *tg* in thyrocytes**

t-SNE plots depicting the thyrocytes clusters (above) along with expression of *pax2a* and *tg* (below).

### Supplementary Table Legends

**Table 1: Marker genes for each cluster of the zebrafish thyroid gland atlas (.xls)**

A table listing the marker genes for the seven clusters identified in the zebrafish thyroid gland atlas. The tables also lists the fold-change and adjusted p-value for each gene.

**Table 2: Connectome for the zebrafish thyroid gland atlas (.xls)**

A table listing the ligands expressed in different cell-types of the thyroid gland and their corresponding receptors expressed on the thyrocytes.

**Table 3: Differential gene expression analysis between the two sub-populations of the thyrocytes (.xls)**

A table listing the differentially expressed genes between the ‘Cluster-Blue’ and ‘Cluster-Red’ subpopulations of the thyrocytes. The table lists fold-change and p-value for each gene. Genes showing a significant difference (p-value < 0.05) in gene expression are listed.

**Table 4: Genes displaying transcriptional heterogeneity within the thyrocyte population (.xls)**

A table listing the genetic entropy, or transcriptional heterogeneity, within the thyrocyte population. Entropy is a measure of the degree of uncertainty in the expression of a gene. The table lists mean expression, entropy measure and p-value for each gene. Genes showing a significant entropy (p-value < 0.05) are listed.

### Supplementary Movie Legend

**Movie 1: Time-lapse of *pax2a*$^{mKO2}$ expression during embryonic development**

A time-lapse video from confocal imaging of the *pax2a*$^{mKO2}$; *Tg(tg:nls-EGFP)* zebrafish embryo from 36 hpf to 55 hpf. Live imaging reveals expression of mKO2 in anatomical structures known to express *pax2a*. Moreover, co-expression of mKO2 and GFP can be observed in the developing thyroid gland.
